# Excitable RhoA dynamics drive pulsed contractions in the early *C. elegans* embryo

**DOI:** 10.1101/076356

**Authors:** Jonathan B. Michaux, François B. Robin, William M. McFadden, Edwin M. Munro

**Author notes:** Equal contributions (Jonathan B. Michaux, François B. Robin).

## Abstract

Pulsed actomyosin contractility underlies diverse modes of tissue morphogenesis, but the underlying mechanisms remain poorly understood. Here, we combine quantitative imaging with genetic perturbations to identify a core mechanism for pulsed contractility in early *C. elegans* embryos. We show that pulsed accumulation of actomyosin is governed by local control of assembly and disassembly downstream of RhoA. Pulsed activation and inactivation of RhoA precede, respectively, accumulation and disappearance of actomyosin, and persist in the nearly complete absence of Myosin II. We find that fast positive feedback on RhoA activation drives pulse initiation, while F-actin dependent accumulation of the RhoA GTPase activating proteins (GAPs) RGA-3/4 provides delayed negative feedback to terminate each pulse. An experimentally constrained mathematical model confirms that in principle these feedbacks are sufficient to generate locally excitable RhoA dynamics. We propose that excitable RhoA dynamics are a common driver for pulsed contractility that can be differently tuned or coupled to actomyosin dynamics to produce a diversity of morphogenetic outcomes.

## Introduction

Pulsed contractility is a widespread mode of actomyosin contractility expressed by many non-muscle cells in which transient accumulations of F-actin and Myosin II accompany local contractions of the cell surface. Pulsed contractions were first identified in the polarizing *C. elegans* zygote (Munro et al., 2004), and have now been documented in a wide variety of embryonic and extra-embryonic epithelia (He et al., 2010; Blanchard et al., 2010; Rauzi et al., 2010; David et al., 2010; Martin et al., 2009; Solon et al., 2009) and mesenchymal cells (Kim and Davidson, 2011). A similar phenomenon known as cell shape oscillations have been observed in many cultured cells (Sedzinski et al., 2011; Salbreux et al., 2007; Kapustina et al., 2008). Pulsed contractions produce transient shape changes that can be biased or rectified in different ways to produce distinct morphogenetic outcomes such as tissue invagination (Martin et al., 2009), tissue elongation (Rauzi et al., 2010; Levayer and Lecuit, 2013; He et al., 2010), epithelial tissue closure (Solon et al., 2009) and wound healing (Razzell et al., 2014). During embryonic development, pulsed contractions may represent an adaptation to accommodate rapid cell and tissue deformations while maintaining overall tissue integrity (Vasquez et al., 2014). In other contexts, such as in many cultured cells, shape oscillations may represent an aberrant behavior that manifests when cells lose normal adhesion to their substrates (Salbreux et al., 2007; Paluch et al., 2005), or when microtubules are depolymerized (Kapustina et al., 2013; 2008; Rankin and Wordeman, 2010; Werner et al., 2007; Piekny and Glotzer, 2008; Bornens et al., 1989) or when contractile tension is very high during cytokinesis (Sedzinski et al., 2011).

Despite the widespread occurrence of pulsed contractions and increasing evidence for their functional relevance, the mechanisms that initiate and terminate pulsed contractions remain poorly understood. From a dynamical perspective, pulsed contractions represent a form of excitable behavior, exemplified by action potentials in neuronal cells (Izhikevich, 2007) or pulses of intracellular calcium release observed in many cell types (Goldbeter, 1996). Theoretical studies highlight two key ingredients for excitability: positive feedback to drive a rapid upswing in activity, and delayed negative feedback to bring it back down again. A key challenge is to identify the specific modes of positive and negative feedback that drive pulsed contractions.

Multiple forms of positive feedback could contribute to initiating pulsed contractions. For example, local actomyosin-based contraction could promote further accumulation of actomyosin through mechanosensitive motor-filament binding (He et al., 2010; Fernandez-Gonzalez et al., 2009; Ren et al., 2009; Effler et al., 2006; Schiffhauer et al., 2016), by enhancing actin filament assembly and/or stability (Hayakawa et al., 2011; La Cruz and Gardel, 2015), or by transporting and concentrating actomyosin and/or its upstream activators (Munjal et al., 2015; Dierkes et al., 2014). Alternatively, dynamic clustering of F-actin and/or Myosin II by scaffolding proteins such as Anillin could promote Myosin II recruitment and focal contraction (Maddox et al., 2005). Finally, autocatalytic activation of upstream regulators such as RhoA could drive local excitation, independent of, or in addition to, myosin-based tension or network contraction (Zhang and Glotzer, 2015; Munjal et al., 2015; Bement et al., 2015; Graessl et al., 2017). Similarly, multiple forms of delayed negative feedback could contribute to terminating pulses, including progressive buildup of steric or elastic resistance to further contraction (Dierkes et al., 2014), or contraction-mediated disassembly, or delayed recruitment of disassembly factors or inhibitors of Myosin II or RhoA (Munjal et al., 2015; Kasza and Zallen, 2011; Bement et al., 2015; Graessl et al., 2017).

Here we combine quantitative imaging with experimental manipulations and mathematical modeling to identify the dynamical basis for pulsed contractility in the early *C. elegans* embryo. Using single-molecule imaging and particle tracking analysis, we provide definite evidence that the initiation of pulsed contractions does not involve or require local redistribution of actomyosin or its upstream activators. Instead, pulsed contractions are driven by local pulses of RhoA activity, which feed forward to control local accumulation of downstream targets F-actin, Myosin II and Anillin. We present evidence that pulsed accumulation of RhoA is governed by locally excitable RhoA dynamics: local autocatalytic activation of RhoA drives the rapid upswing of RhoA activity during pulse initiation, while F-actin-dependent recruitment of the redundantly acting RhoA GTPase activating proteins (GAPs) RGA-3/4 provide delayed negative feedback to terminate the pulse. A minimal model, sharply constrained by our experimental data, suggests that this combination of feedbacks is sufficient to generate locally excitable or oscillatory RhoA dynamics and to explain quantitatively the temporal dynamics of RhoA activation and RGA-3/4 accumulation during each pulse. We propose that excitable RhoA dynamics define a core mechanism for pulsed contractility and suggest that this mechanism may be tuned or filtered through downstream effectors to control the size or spacing or lifetime of pulsed contractions.

Here we combine quantitative imaging with experimental manipulations and mathematical modeling to identify the dynamical basis for pulsed contractility in the early *C. elegans* embryo. Using single-molecule imaging and particle tracking analysis, we provide definite evidence that the initiation of pulsed contractions does not involve or require local redistribution of actomyosin or its upstream activators. Instead, pulsed contractions are driven by local pulses of RhoA activity, which feed forward to control local accumulation of downstream targets F-actin, Myosin II and Anillin. We present evidence that pulsed accumulation of RhoA is governed by locally excitable RhoA dynamics: local autocatalytic activation of RhoA drives the rapid upswing of RhoA activity during pulse initiation, while F-actin-dependent recruitment of the redundantly acting RhoA GTPase activating proteins (GAPs) RGA-3/4 provide delayed negative feedback to terminate the pulse. A minimal model, sharply constrained by our experimental data, suggests that this combination of feedbacks is sufficient to generate locally excitable or oscillatory RhoA dynamics and to explain quantitatively the temporal dynamics of RhoA activation and RGA-3/4 accumulation during each pulse. We propose that excitable RhoA dynamics define a core mechanism for pulsed contractility and suggest that this mechanism may be tuned or filtered through downstream effectors to control the size or spacing or lifetime of pulsed contractions.

## Results

Pulsed contractions were originally described in *C. elegans* during interphase in the polarizing zygote P0 (Movie S1, top; Figure 1 A-C, individual pulses indicated by white arrowheads in Figure 1B; (Munro et al., 2004)). In these cells, pulsed contractions are associated with transient deep invaginations of the cell surface (magenta arrows in Figure 1A,B); this makes it more difficult to quantify local changes in density of cortical factors during individual pulses, because these measurements could be confounded by movements of the cortex in/out of the plane of focus. Therefore, we focused on pulsed contractions that occur at the two-cell stage in the anterior blastomere known as AB (Figure 1D-F, individual pulses indicated by white arrowheads in Figure 1E). As in P0, pulsed contractions in AB involve transient accumulations of F-actin and Myosin II; they are associated with transient local contractions of the actomyosin cortex (Figure 1F), but they are not associated with pronounced invaginations of the cell surface.

**Figure 1.**
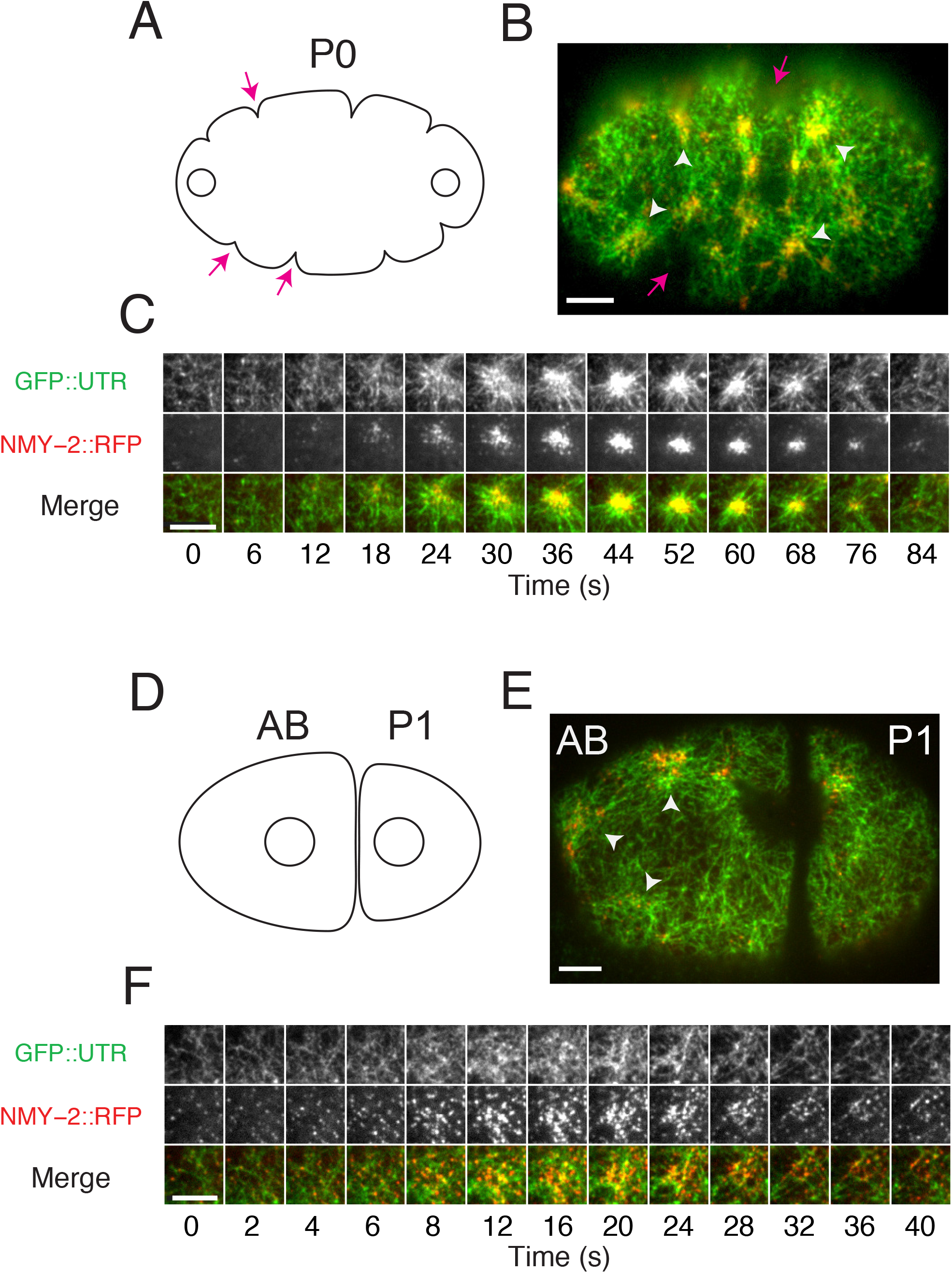
Actomyosin pulses in 1 and 2-cell stage embryos. (A) Schematic view of the zygote P0 during early interphase. Open circles represent the two pronuclei. (B) Micrograph of P0 in early interphase expressing GFP::UTR and NMY-2::RFP. In (A) and (B), white arrowheads indicate individual pulses and magenta arrows indicate membrane invaginations (ruffles). (C) Time evolution of a single pulse in P0. The time delay between frames is 6s for the first 6 frames, and 8s thereafter. (D) Schematic of an embryo at the early 2-cell stage, showing the anterior blastomere AB and the posterior blastomere P1. Open circles represent the interphase nuclei. (E) Micrograph of an early two-cell stage embryo expressing GFP::UTR and NMY-2::RFP. White arrowheads indicate individual pulses (F) Time evolution of a single pulse. The time delay between frames is 2s for the first 5 frames, and 4s thereafter. Scale bars = 5µm in B,C,E,F.

**Table 1.**
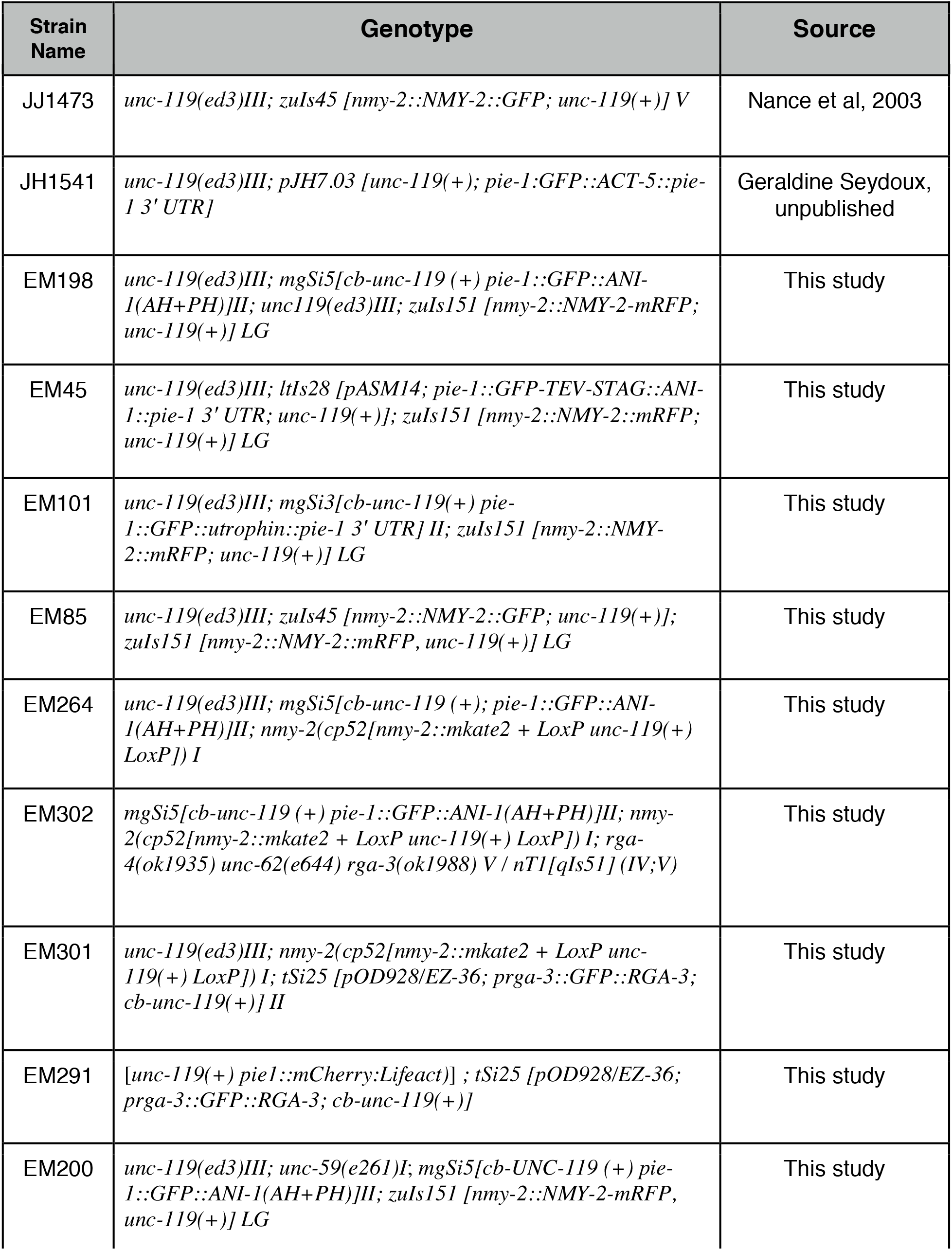
List of strains used in this study

### Single-molecule analysis of actomyosin dynamics during pulsed contractions

As a key step towards distinguishing different mechanisms for pulsed contractions, we used single-molecule imaging and single-particle tracking analysis to quantify the relative contributions of local turnover and redistribution to changes in F-actin and Myosin II density during individual pulses. As described previously (Robin et al., 2014), we used RNAi against GFP to obtain embryos expressing single molecule levels of Actin::GFP or of the non-muscle myosin heavy chain fused to GFP (NMY-2::GFP) over the endogenous proteins (Figure 2A, Movies S2,4). We combined near-total internal reflection fluorescence (TIRF) imaging with single-molecule detection and tracking to measure the appearance, motion and disappearance of single-molecule speckles (Figure 2A-C, (Robin et al., 2014)). We assumed, with others (Watanabe and Mitchison, 2002; Vallotton et al., 2004), that single molecule appearance and disappearance events report directly on local rates of filament assembly and disassembly. We have shown previously that rates of turnover measured by single-molecule tracking agree well with those measured from single-molecule data by fitting kinetic models to photobleaching curves ((Robin et al., 2014), Materials and Methods).

**Figure 2.**
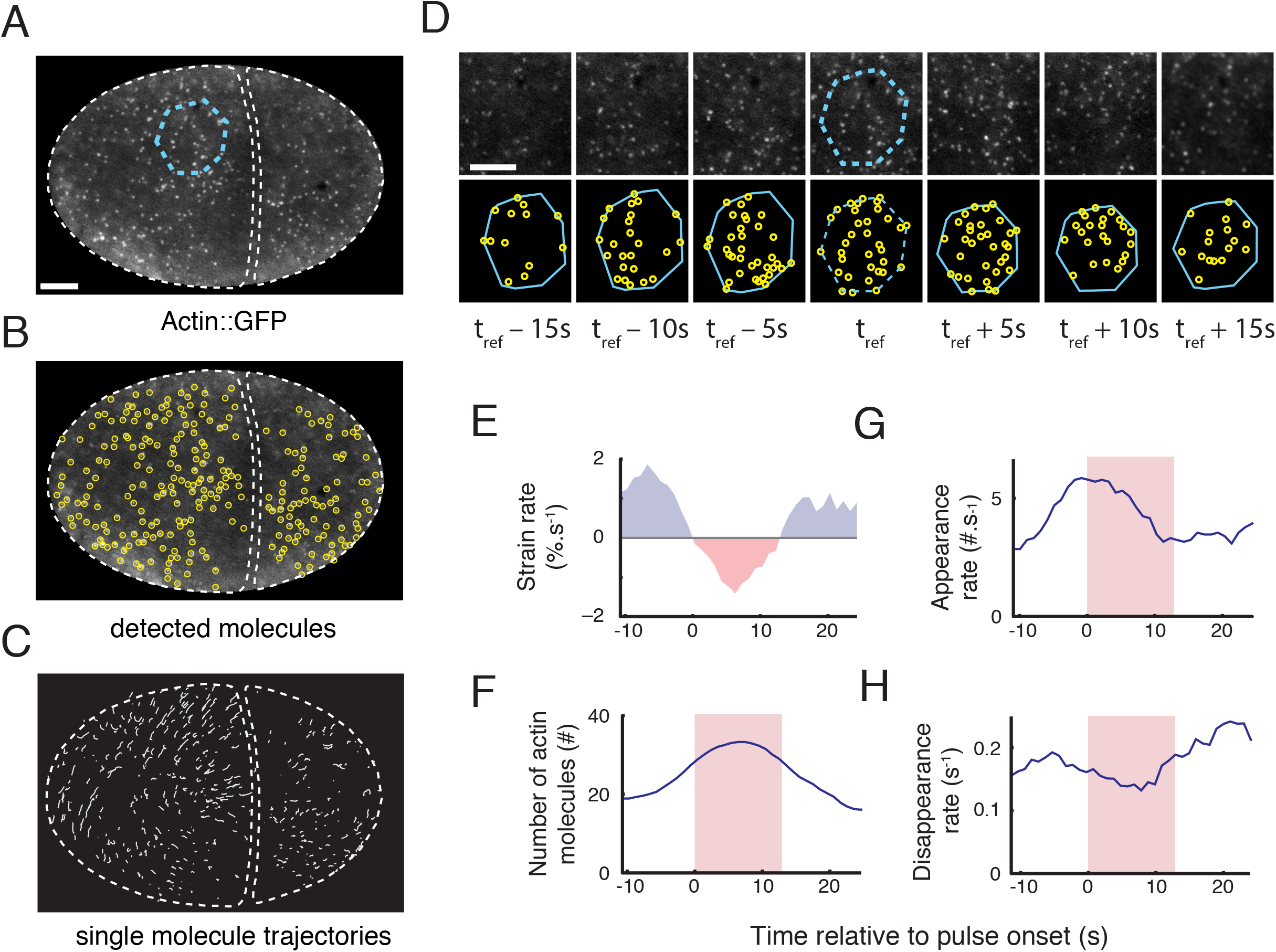
Single-molecule analysis of actin network assembly, disassembly and deformation during individual pulsed contractions. (A) One frame of a time lapse sequence taken from an embryo expressing low levels of Actin::GFP. A patch of cortex undergoing a pulse is identified from the time lapse sequence, and outlined in cyan. (B) Automatic particle detection of single-molecules from the image in (A). (C) Trajectories of the molecules displayed in (B) that were tracked for longer than 2s. (D) A polygonal region of interest identified at time t = t_ref_, in (A) (dashed cyan polygon) is propagated forward and backward in time using the trajectories of tracked particles (see Materials and Methods and Movies S2, S3). (E-H) Simultaneous measurements of single molecule dynamics and patch deformation over time. (E) Strain rate, measured using the particle-based method (see Materials and methods). (F) Number of actin molecules in the patch. (G-H) Appearance rates (G) and disappearance rates (H) of actin molecules. Red shading in (F-H) indicates the time period in which the cortex is contracting locally (strain rate <0). Scale bars = 5µm in A; 3µm in D.

We then devised new methods to measure simultaneously: (a) single molecule appearance rates, disappearance rates and densities, and (b) local rates of cortical deformation, on a moving and deforming patch of cortex during individual pulsed contractions (Figure S1; see Materials and Methods for details). Briefly, we identified a reference frame for each pulse near the onset of contraction; within that frame, we identified a polygonal region of interest containing the contracting patch (dashed blue polygon in Figure 2A; hereafter “the patch”); we propagated the patch forward and backwards in time by extrapolating the displacements of tracked particles on or near its boundary (Figure 2D, Figure S1, Movie S3). We then measured local deformation of the patch as frame-to frame changes in patch area, or by estimating a local strain rate from frame-to-frame displacements of the individual particles, with similar results in both cases (Figure 2E; Figure S2A-C; Materials and Methods). At the same time, we measured the number of molecules and single-molecule appearance and disappearance rates within the patch over time (Figure 2F-H). Finally, we aligned single-molecule measurements with respect to the onset or termination of individual contractions to produce a dynamical signature of actin assembly, disassembly and deformation over the lifetime of a pulse (Figure 3A-D; Figure S2D-F). These measurements allowed us to distinguish, cleanly, changes in single-molecule densities due to local assembly and disassembly from those due to local contraction (or expansion) of the cortical patch.

**Figure 3.**
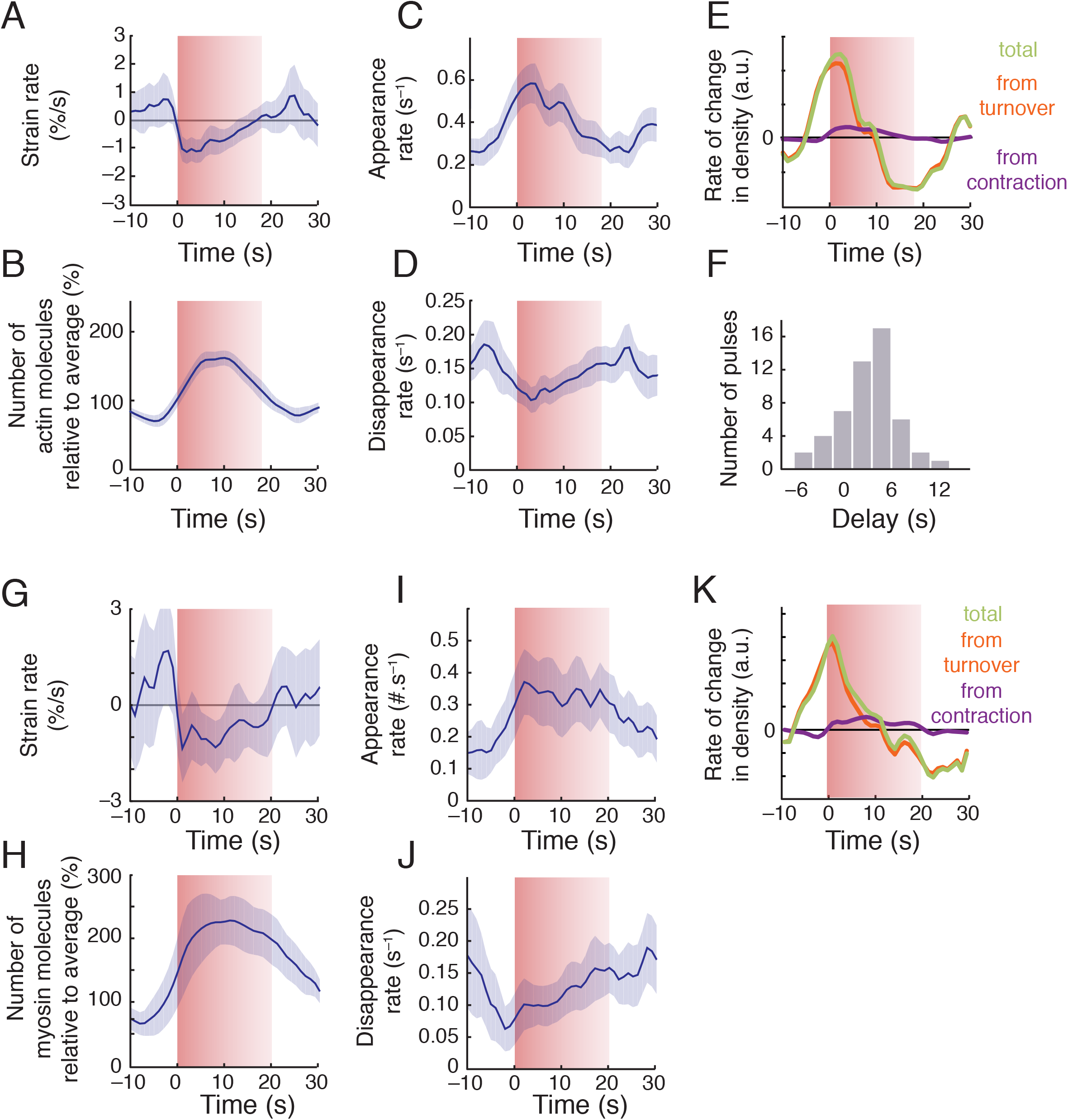
Spatiotemporal modulation of assembly and disassembly drives transient accumulation of F-actin and Myosin II during pulsed contractions. (A-D) Data from individual pulses, aligned with respect to the onset of contraction and then averaged to display particle-based strain rate (A), numbers of actin molecules (scaled by the average number for each pulse) (B) appearance rate (C), and disappearance rate (D) versus time. (E) Total rate of change in actin density (green) and the individual contributions to rate of change from turnover (red) and surface contraction (purple). (F) Distribution of time delays between the initiation of contraction and actin accumulation. Data in (A-F) were averaged over 42 pulses, collected in 8 embryos. (G-J) Average myosin dynamics synchronized with respect to time at which myosin density reached peak levels during a pulse, displaying particle-based strain rate (G), number of molecules (scaled to the average number for each pulse) (H), appearance rate (I) and disappearance rate (J) versus time. (K) Total rate of change in myosin density (green) and the individual contributions to rate of change from turnover (red) and surface contraction (purple). Data in (G-K) were averaged over 30 pulses, collected in 5 embryos. Error bars: 95% confidence interval.

If pulses are initiated by positive feedback in which local contraction concentrates actomyosin and/or its upstream regulators, then the onset of actomyosin accumulation should coincide with the onset of contraction. Contradicting this expectation, we found that, on average, Actin:GFP began to accumulate ~5s before the onset of contraction (Figure 3B,F; Figure S2A-C), during a period of time in which the cortex was locally expanding (Figure 3A). Approximately 30% of the total increase in Actin::GFP single molecule density measured during a pulse occurred before the onset of contraction (Figure S2A-C). This initial accumulation was due entirely to a net imbalance of assembly and disassembly (Figure 3E): Before the onset of contraction, assembly rates increased (Figure 3C) and disassembly rates decreased (Figure 3D), leading to a sharp increase in the net rate of single molecule accumulation that peaked at the onset of contraction (Figure 3E). During the contraction phase itself, the rate of change in single-molecule densities was determined almost entirely by a net imbalance of assembly/disassembly, with a very minor (< 6%) contribution from contraction itself (Figure 3E). Assembly rates decreased steadily, and disassembly rates increased steadily, such that a transition from net assembly to net disassembly (and from increasing density to decreasing density) occurred ~7s sec after the onset of contraction (Figure 3E). We obtained very similar results in embryos depleted of ARX-2, an essential subunit of the ARP2/3 complex (Figure S3), suggesting that our results are not strongly biased by selective incorporation of Actin::GFP into branched vs. unbranched F-actin (Chen et al., 2012).

Single-molecule analysis of GFP-tagged Myosin II (NMY-2::GFP) revealed local assembly/disassembly dynamics that were strikingly similar to those measured for GFP::Actin. (Figure 3G-K). On average, the density of single molecules of NMY-2::GFP began to increase ~6s before the onset of contraction during a period of local cortical expansion (Figure 3H), and approximately 50% of this increase occurred before the onset of contraction. As observed for GFP::Actin, the upswing in Myosin II before the onset of a contraction was associated with both a sharp increase in appearance rates and a sharp decrease in disappearance rates (Figure 3I,J); the net rate of increase peaked at the onset of contraction, and during the contraction phase, the rate of change in single-molecule densities was determined largely by a net imbalance of assembly/disassembly, with a minor contribution from contraction (Figure 3K).

In summary, we find that changes in actomyosin density during pulsed contractions are governed primarily by dynamic local imbalance of F-actin and Myosin II appearance and disappearance rates. A large fraction of the increase in F-actin and Myosin II density during each pulse occurs before the onset of contraction, and local contraction accounts for only a minor fraction of the subsequent density increase during the contraction phase itself. We conclude that changes in actomyosin density during pulsed contractions are governed primarily by dynamic modulation of assembly and disassembly, not by local clustering of these factors or by dynamical coupling of contraction and advection.

### Pulsed activation of RhoA drives the pulsed accumulation of F-actin and Myosin II

The observation that F-actin and Myosin II accumulate with very similar kinetics during pulsed contractions suggests that their accumulation is driven by a common upstream regulator. An obvious candidate is the small GTPase RhoA (encoded by *rho-1* in *C. elegans*), which recruits and/or activates downstream effectors including formins, Rho Kinase (ROCK) and Anillin to control F-actin assembly and Myosin II activation in a variety of cell types (Jaffe and Hall, 2005; Piekny and Glotzer, 2008). RhoA activity is required for pulsed contractions in P0 (Motegi:2006hi; Schonegg:2007if; Tse et al., 2012), and a biosensor for active RhoA derived from the RhoA binding domain of Anillin (hereafter GFP::AHPH) accumulates locally during pulsed contractions in the zygote (Tse et al., 2012; Nishikawa et al., 2017).

To determine the relative timings of RhoA activation and actomyosin accumulation during pulsed contractions, we used a strain co-expressing GFP::AHPH (Tse et al., 2012) with an RFP-tagged version of the myosin heavy chain (NMY-2::RFP) to co-monitor RhoA activity and Myosin II accumulation during individual pulses in AB. We observed a striking correlation between pulsed accumulation of GFP::AHPH and NMY-2::RFP during individual pulsed contractions (Figure 4A-C, Movie S5). GFP::AHPH accumulated rapidly within a broad domain that prefigured the initial accumulation of NMY-2::RFP, reached a peak near the onset of visible contraction, and then began to disappear before NMY-2::RFP (Figure 4B,C). NMY-2::RFP accumulated as discrete particles that increased in number and size, while contracting together into a smaller and tighter central domain. The initial accumulation of GFP::AHPH was diffuse, but over time, a fraction of the total GFP:AHPH became enriched within brighter punctae that colocalized with NMY-2::RFP (yellow arrows in Figure 4B). During the falling phase of each pulse, the diffuse pool of GFP::AHPH disappeared rapidly, while a smaller and more persistent fraction of GFP::AHPH remained co-localized with NMY-2::RFP particles (yellow arrows in Figure 4B).

**Figure 4.**
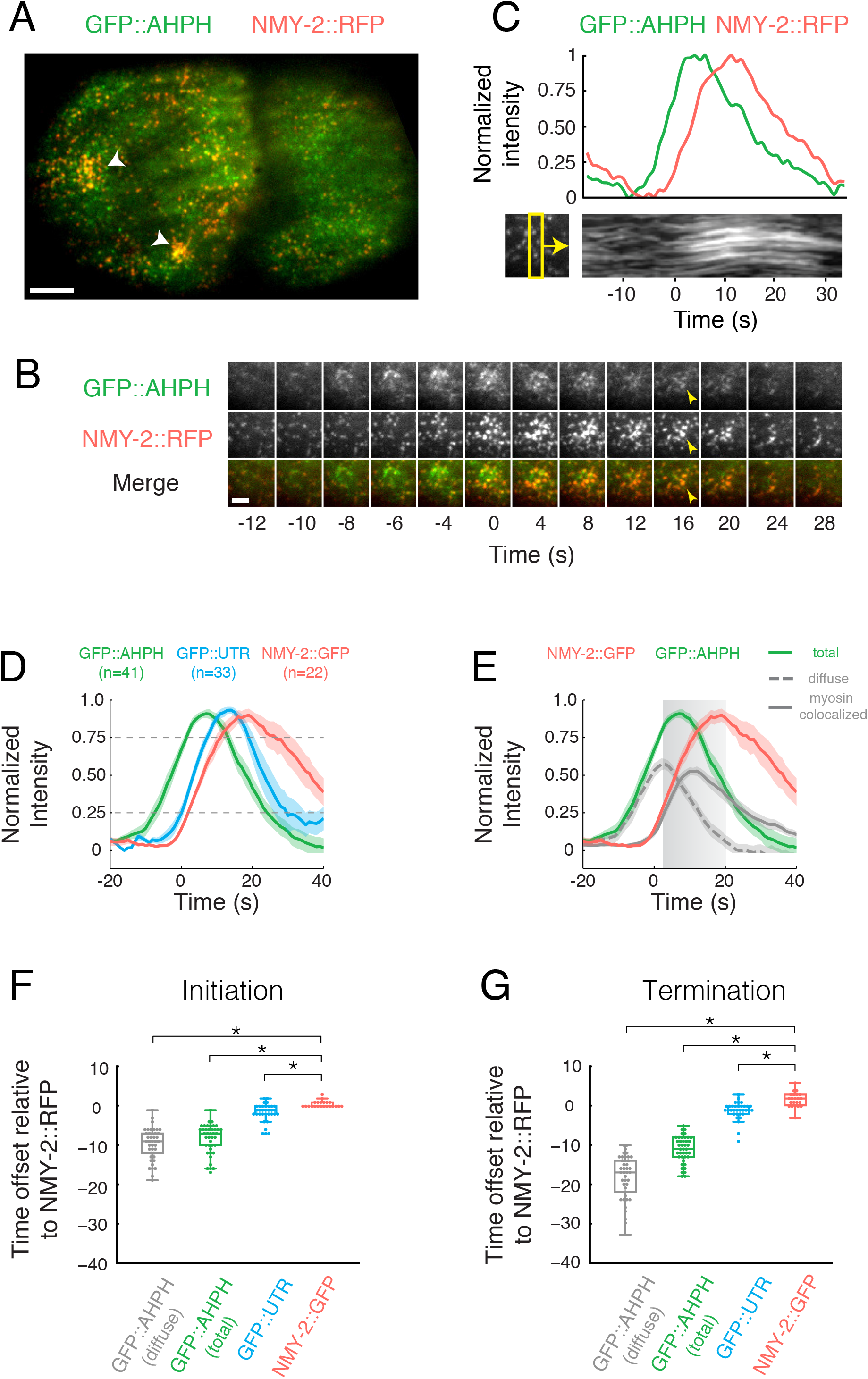
Local pulses of RhoA activation underlie pulsed accumulation and disappearance of F-actin and Myosin II. (A) Micrograph of a 2-cell stage embryo expressing GFP::AHPH as a reporter for RhoA activity, and NMY-2::RFP. (B) Temporal dynamics of GFP::AHPH and NMY-2::RFP accumulation during a single pulse. The time between frames is 2s for the first five frames, and 4s thereafter. (C) Top: Normalized fluorescence intensities of GFP::AHPH, and NMY-2::RFP. Bottom: A kymograph showing that local contraction (concerted movements of myosin puncta) begins after the accumulation of GFP::AHPH. The yellow box below left indicates the region used to generate the kymograph. (D) Comparison of averaged normalized fluorescence intensities vs. time for active RhoA (GFP::AHPH), Myosin (NMY-2::GFP) and F-actin (GFP::UTR) from two-color data, co-aligned with respect to a common reference signal (NMY-2:RFP). Data were co-aligned with respect to the time at which NMY-2::RFP reaches25% threshold. Hued regions report 95% confidence intervals. (E) Decomposition of the GFP::AHPH signal into a diffuse pool (gray, dashed line) and myosin-colocalized pool (gray, solid line) (see Figure S3 and Materials and Methods for details). Data for NMY-2::RFP and total GFP::AHPH are the same as in panel D. (F) Distribution of the delays between the onset of accumulation of NMY-2:RFP, and the onset of accumulation of GFP::AHPH, NMY-2::GFP and GFP::UTR. Onset of accumulation was measured as the time at which normalized probe intensity rose above 25% of its maximal level. (G) Distribution of the delays between the onset of disappearance of NMY-2:RFP, and the onset of disappearance of GFP::AHPH, NMY-2::GFP and GFP::UTR. Onset of disappearance was measured as the time at which normalized probe intensity fell below 75% of its maximal level. In (F,G) box plots, the central mark represents the median, the box indicates the 25th and 75th percentile, the whiskers mark the minimum and maximum values and the “+” symbol represents outliers. Scale bars = 5µm in A; 2µm in B.

To quantify these observations, we aligned data for multiple pulses from embryos co-expressing NMY-2::RFP and GFP::AHPH as follows (see Figure S3; Materials and Methods). For each pulse, we smoothed and thresholded the NMY-2::RFP signal to identify a region of interest (ROI) containing high levels of NMY-2::RFP just before the onset of contraction (Figure S3A; Materials and Methods). We propagated this ROI forward and backwards in time (Figure S3A). We measured the mean intensities of the NMY-2::RFP and GFP::AHPH signals within the ROI before, during and after the pulse, and normalized these data with respect to the minimum (pre-contraction) and maximum intensities measured during this interval (Figure S3B). Then we aligned data for multiple pulses with respect to the time point at which NMY-2::RFP reached 25% of its maximum intensity (Figure S3C, Figure 4D). To distinguish different GFP::AHPH fractions, we thresholded the raw NMY-2::RFP signal to create a binary mask that distinguishes regions of high and low Myosin II intensity within the ROI (Figure S3D). Then we used this mask to decompose the normalized AHPH::GFP signal into “diffuse” and “myosin-colocalized” fractions (Figure S3E-F).

These aligned data confirm that sharp increases and decreases in total GFP::AHPH intensity precede, respectively, the appearance and disappearance of NMY-2::RFP (Figure 4D). On average, total GFP::AHPH reached 25% of its maximum intensity 8.6 +/-3.9 seconds before NMY-2::RFP (Figure 4F), and fell below 75% of its maximum intensity 11.1 +/-3.5 seconds before NMY-2::RFP (Figure 4G). Consistent with direct observations, the mean intensity of the diffuse fraction rose in phase with, but peaked earlier than the total GFP::AHPH signal, while the mean intensity of the myosin-colocalized fraction rose in phase with NMY-2::RFP and peaked later than the total GFP::AHPH signal (Figure 4E-G).

We used the same approach to align data for embryos co-expressing NMY-2::RFP and the F-actin binding domain of Utrophin fused to GFP (GFP::UTR); a marker for F-actin (Burkel et al., 2007; Tse et al., 2012). Using NMY-2::RFP as the common reference to co-align data for NMY-2::RFP, GFP::AHPH, and GFP::UTR, we found that, like Myosin II, F-actin accumulates and dissipates during pulsed contractions with a significant delay relative to GFP::AHPH (Figure 4D,F,G). Thus local activation and inactivation of RhoA precedes the accumulation and disappearance of its downstream targets.

Finally, we used the time points at which NMY-2::RFP intensities (Figure 4D) and single molecule densities of Myosin::GFP (Figure 3H) reached 25% of their peak values to align the time course of GFP::AHPH accumulation with respect to the onset of contraction as measured by single molecule imaging. This analysis shows that the GFP::AHPH signal peaks just before (diffuse signal) or just after (total signal) the onset of contraction (indicated by grey box in Figure 4E). Thus local concentration of active RhoA by advection-contraction (Munjal et al., 2015) cannot explain the rising phase of RhoA activation in *C. elegans* embryos. Moreover, the diffuse GFP::AHPH signal peaks and begins to fall at a point where F-actin and Myosin II disappearance rates are at a minimum (Figure 3D,J); thus factors other than cortical actomyosin disassembly drive the disappearance of active RhoA at the end of a pulse.

We used the same two-color image analysis to characterize co-accumulation of GFP::AHPH, NMY-2::RFP and GFP::UTR during pulsed contractions in the zygote P0. To remove the potentially confounding effects of large scale cortical flows that occur during zygote polarization, we performed these measurements in embryos depleted of the centrosomal factor SPD-5 (Munro et al., 2004; Hamill et al., 2002), which exhibit pulsed contractions but lack cortical flows. In *spd-5(RNAi)* P0 zygotes, as in wild type AB cells, increases in local GFP::AHPH intensity preceded the rise of NMY-2::RFP (Figure S4B-D) and to a lesser extent GFP::UTR (Figure S4D). As in AB, the initial accumulation of GFP::AHPH was broad and diffuse while a more persistent pool of punctate GFP::AHPH co-localized with NMY-2::RFP within a more central region of the initial domain (yellow arrows in Figure S4B). The duration of GFP::AHPH and NMY-2::RFP accumulation was significantly longer in *spd-5(RNAi)* P0 zygotes than in wild type AB cells. However, decomposition of the GFP::AHPH signal into “diffuse” and “myosin colocalized” fractions (Figure S3F, S4E) revealed that the diffuse GFP::AHPH signal rose and fell with very similar temporal dynamics in AB and P0, while the myosin co-localized signal is far more persistent in P0 than in AB (compare Figure S3E and F).

### Pulsed activation of RhoA does not require Myosin II

The observation that active RhoA peaks before the onset of contraction rules out models in which local contraction concentrates RhoA, or its upstream activators, to initiate pulses (Munjal et al., 2015). However, Myosin II could act in other ways to promote or shape pulsed activation of RhoA. To test this possibility, we used RNAi to deplete NMY-2 in a strain co-expressing transgenic GFP::AHPH and NMY-2 tagged with mKate2 at the endogenous locus by CRISPR-mediated homologous recombination (a kind gift of Dan Dickinson). We found that local pulses of GFP::AHPH accumulation persisted in zygotes almost completely depleted of NMY-2, such that fewer than two NMY-2::mKate2-containing particles could be detected during a given pulse (Figure 5A,B Movie S6). Focal accumulations of GFP::AHPH in *nmy-2(RNAi)* zygotes were comparable in size and spacing to those observed in *spd-5 (RNAi)* zygotes. However the GFP::AHPH signal was entirely diffuse; the punctate signal that colocalizes with Myosin II in wild type embryos was absent; and the mean time course for GFP::AHPH accumulation and disappearance was very similar to that measured for the diffuse pool of GFP::AHPH both in wild type AB cells and in *spd-5(RNAi)* zygotes (Figure 5C). Assuming the diffuse AHPH signal reflects active RhoA, these data reveal the existence of a mechanism for locally pulsatile activation of RhoA, with similar kinetics in P0 and AB cells, that does not require the presence or activity of Myosin II. Whether the more persistent pool of myosin-colocalized AHPH reflects myosin-dependent recruitment and/or stabilization of active RhoA, or binding of the Anillin AHPH domain to other cortical factors, remains unclear (see Discussion).

**Figure 5.**
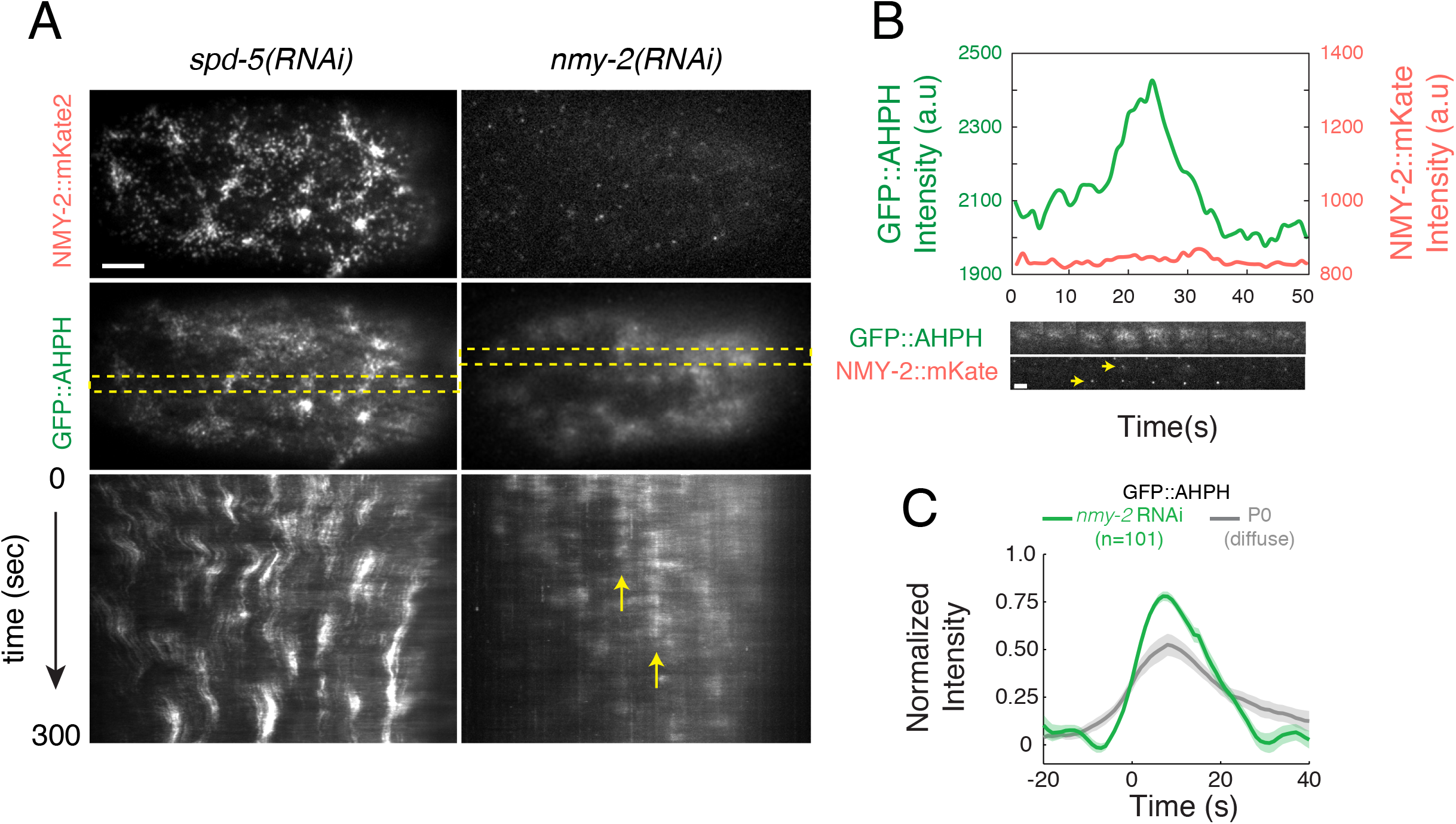
Myosin II is not required for the pulsed activation of RhoA. (A) Comparison of pulse dynamics in zygotes expressing GFP::AHPH and NMY-2::mKATE and treated with either *spd-5(RNAi)* or *nmy-2(RNAi)*. Top panels show myosin localization (NMY-2::mKATE), middle panels show RhoA activity (GFP::AHPH). Bottom panels are kymographs showing GFP::AHPH dynamics over time. Dashed yellow rectangles in middle panel indicate the regions from which the kymographs were made. Vertical yellow arrows indicate a region undergoing repeated pulses. Intensities were scaled identically for *spd-5(RNAi)* and *nmy-2(RNAi)* zygotes. (B) Top: Mean intensities of NMY-2::mKATE (red) and GFP::AHPH (green) vs time for a single pulse in an *nmy-2(RNAi)* zygote. Bottom: sequential snapshots of the region undergoing the pulse showing NMY-2::mKATE (red) and GFP::AHPH (green) distributions. Yellow arrowheads indicate the two particles that can be detected in the region undergoing a pulse. (C) Comparison of total GFP::AHPH during pulses in *nmy-2(RNAi)* zygotes (green curve) with the diffuse pool of GFP::AHPH in *spd-5(RNAi)* zygotes (red curve) and wild type AB cells (blue curve). For each condition, data were aligned with respect to the time at which the normalized signal reached 25% of its maximum value. Data for diffuse AHPH::GFP in *spd-5(RNAi)* zygotes and wild type AB cells are identical to those shown in Figures S4E and 4E respectively. (see Figures 4E, S3D-F and S4E for details). Hued regions report 95% confidence intervals. Scale bars = 5µm in A; 2µm in B.

Another key regulator of pulsed contractions is the scaffold anillin, which binds F-actin, Myosin II, active RhoA, and other cortical factors. The *C. elegans* anillin ANI-1 promotes focal accumulations of Myosin II during polarization and cytokinesis (Maddox et al., 2007; Tse et al., 2011), but whether it does so by promoting RhoA activation, or by recruiting/stabilizing Myosin II downstream of RhoA, or both, remains unclear. To distinguish these possibilities, we first used two color image analysis to examine the relative timing of GFP::AHPH and GFP::ANI-1 accumulation during individual pulses, using NMY-2::RFP as a common reference. In both AB and P0 cells, GFP::ANI-1 accumulated slightly before NMY-2::GFP, and thus with a pronounced delay relative to GFP::AHPH, although the delay was smaller in P0 (Figure 6A-H). In AB cells, the GFP::AHPH signal reached its maximum rate of accumulation before a detectable increase in GFP::ANI-1 (Figure 6C), suggesting that a local increase in ANI-1 is not required for rapid activation of RhoA during pulse initiation.

**Figure 6.**
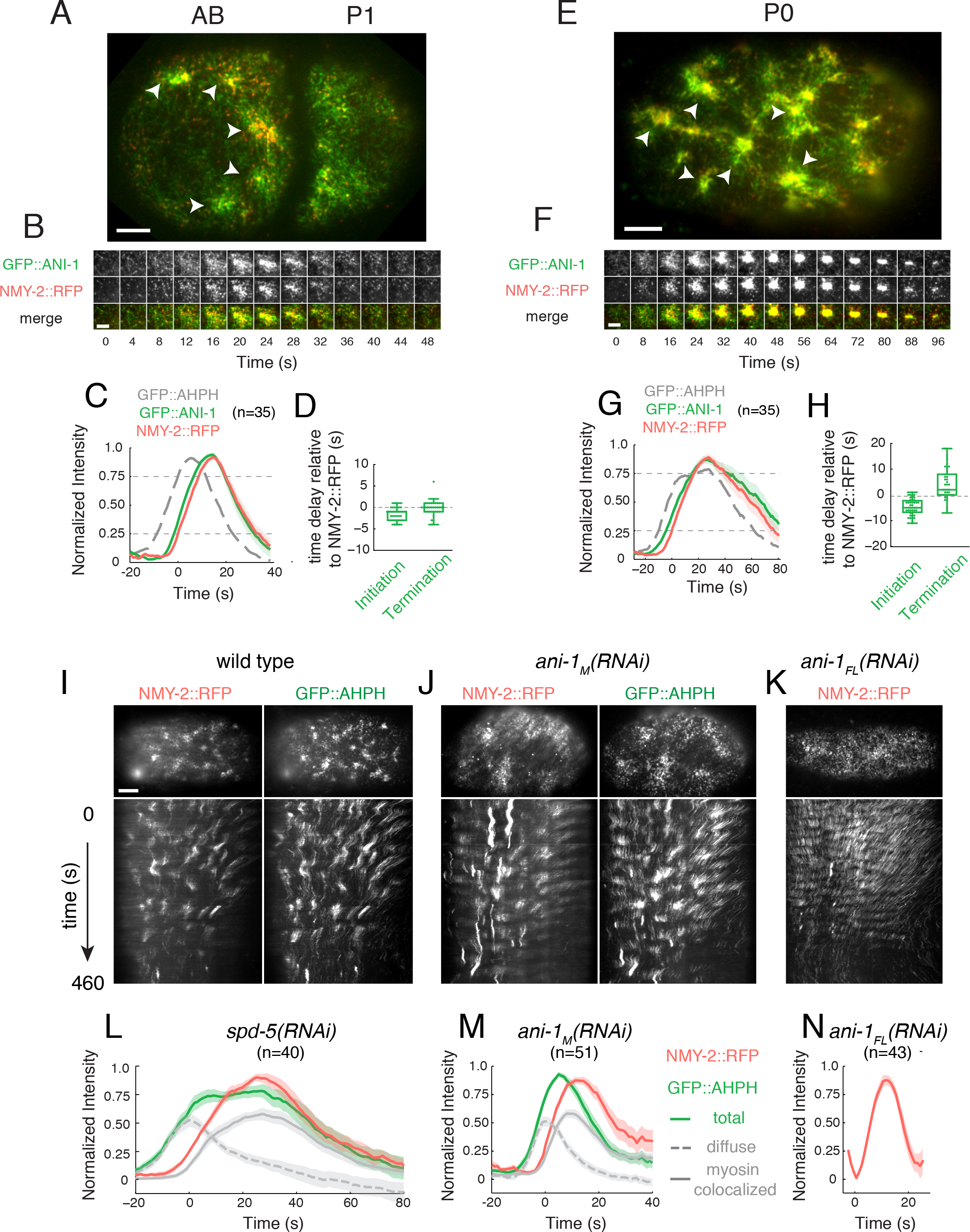
Two-color analysis of Anillin dynamics and contributions to pulsed contractions in one- and two-cell embryos. (A,E) Micrographs of two-cell (A) and one-cell (E) embryos co-expressing GFP::ANI-1 and NMY-2::RFP. White arrowheads indicate individual pulses. (B,F) Expanded views of single pulses illustrating temporal dynamics of GFP::ANI-1 and NMY-2::RFP accumulation. (C,G) Plots of averaged normalized fluorescence intensities for NMY-2::RFP and GFP::ANI-1 from two-color movies, aligned to the time at which NMY-2::RFP reaches 25% threshold. The averaged normalized fluorescence intensity of GFP::AHPH, co-aligned using the NMY-2::RFP signal, is shown for reference. Halos report 95% confidence intervals. (D,H) Distribution of the delays in the onset of appearance and disappearance of GFP::ANI-1 measured relative to NMY-2::RFP. Onset of appearance and disappearance were measured respectively as the time at which the normalized signal rose above 25% or fell below 75% of the maximum value. (I-K) Single frames (top) and kymographs (bottom) comparing NMY-2::RFP and GFP::AHPH dynamics in (I) wild type, (J) *ani-1*_M_(*RNAi*), and (*K*) *ani-1*_FL_*(RNAi)* zygotes. (L,M) Plots of averaged normalized fluorescence intensities of NMY-2::RFP and GFP::AHPH (total, diffuse and myosin-colocalized) for individual pulses from two-color movies in (L) *spd-5(RNAi)* and (M) *ani*-1_M_ *(RNAi)* zygotes, aligned to the time at which NMY-2::RFP reaches 25% threshold. Data in L are identical to those shown in Figure S4E. (N) Plots of averaged normalized fluorescence intensities of NMY-2::RFP for individual pulses in *ani-1*_FL_*(RNAi)* zygotes, aligned to the time at which NMY-2::RFP reaches 25% threshold. Scale bars = 5µm in A,E,I-K; 3µm in B,F.

To test this further, we depleted zygotes of ANI-1 by RNAi in two different ways: RNAi against the full length ani-1 sequence (*ani-1_FL_(RNAi)*) produced strong depletion of both ANI-1 and GFH::AHPH, while RNAi against a shorter N terminal portion of the ani-1 sequence (*ani-1_N_*(*RNAi*)) produced weaker depletion, but without affecting GFP::AHPH expression. In ani-1_FL_(RNAi) zygotes, we still observed pulsed accumulations of NMY-2::RFP, but pulse duration was sharply reduced relative to wild type or *spd-5(RNAi)* zygotes (Fig S5I,K,L,N)), consistent with previous reports (Maddox et al., 2007; Tse et al., 2011). In *ani-1_N_*(*RNAi*) zygotes, the duration of NMY-2::RFP pulses was also sharply reduced, although to a lesser extent than in *ani-1_FL_*(*RNAi*) (Figure 6J,M). The duration of GFP:AHPH pulses was also strongly reduced, due mainly to reduction of the myosin-colocalized signal (Figure 6J,M). In contrast, the diffuse GFP::AHPH signal rose and fell with kinetics that were very similar those observed in *spd-5(RNAi)* zygotes (compare Figure 6L&M). These data argue against an important role for ANI-1 in promoting activation of RhoA during pulse initiation, and instead support a key role for ANI-1 in stabilizing Myosin II downstream of RhoA.

### RhoA feeds back locally to promote its own activity and this is required for pulse initiation

What drives the rapid rise in RhoA activity during pulse initiation? Plotting the rate of change in GFP::AHPH intensity vs intensity during the rising phase of individual pulses in either P0 or AB cells revealed a sharp increase in the rate of RhoA activation with increasing RhoA (Figure 7A). This is consistent with a scenario in which active RhoA feeds back positively to promote further activation of RhoA. However, it could also reflect pulsed activation of RhoA (without feedback) by an upstream activator. To distinguish these possibilities, we used RNAi to progressively deplete embryos of RhoA. If the time course of RhoA activation is dictated by an upstream activator, we should observe pulsed accumulation of GFP::AHPH as long as it remains detectable at the cortex. In contrast, if positive feedback of RhoA onto itself drives pulse initiation, then there should be an abrupt loss of pulsing below a threshold level of RhoA. Consistent with the latter expectation, we observed an abrupt transition from pulsed to non-pulsed RhoA accumulation after ~ 12 hours of feeding (Figure 7B,C, Movie S7). In zygotes that lacked pulsed RhoA accumulation, we could still readily detect robust RhoA-dependent cortical flows (Motegi and Sugimoto, 2006; Schonegg and Hyman, 2006) during polarity establishment (dashed yellow lines in Figure 7B) and localized accumulation of active RhoA prior to cytokinesis (cyan arrowheads in Figure 7B). Together, these observations support the idea that RhoA feeds back positively to amplify its own activation and that sufficiently strong feedback is required to generate local pulses of high RhoA activity.

**Figure 7.**
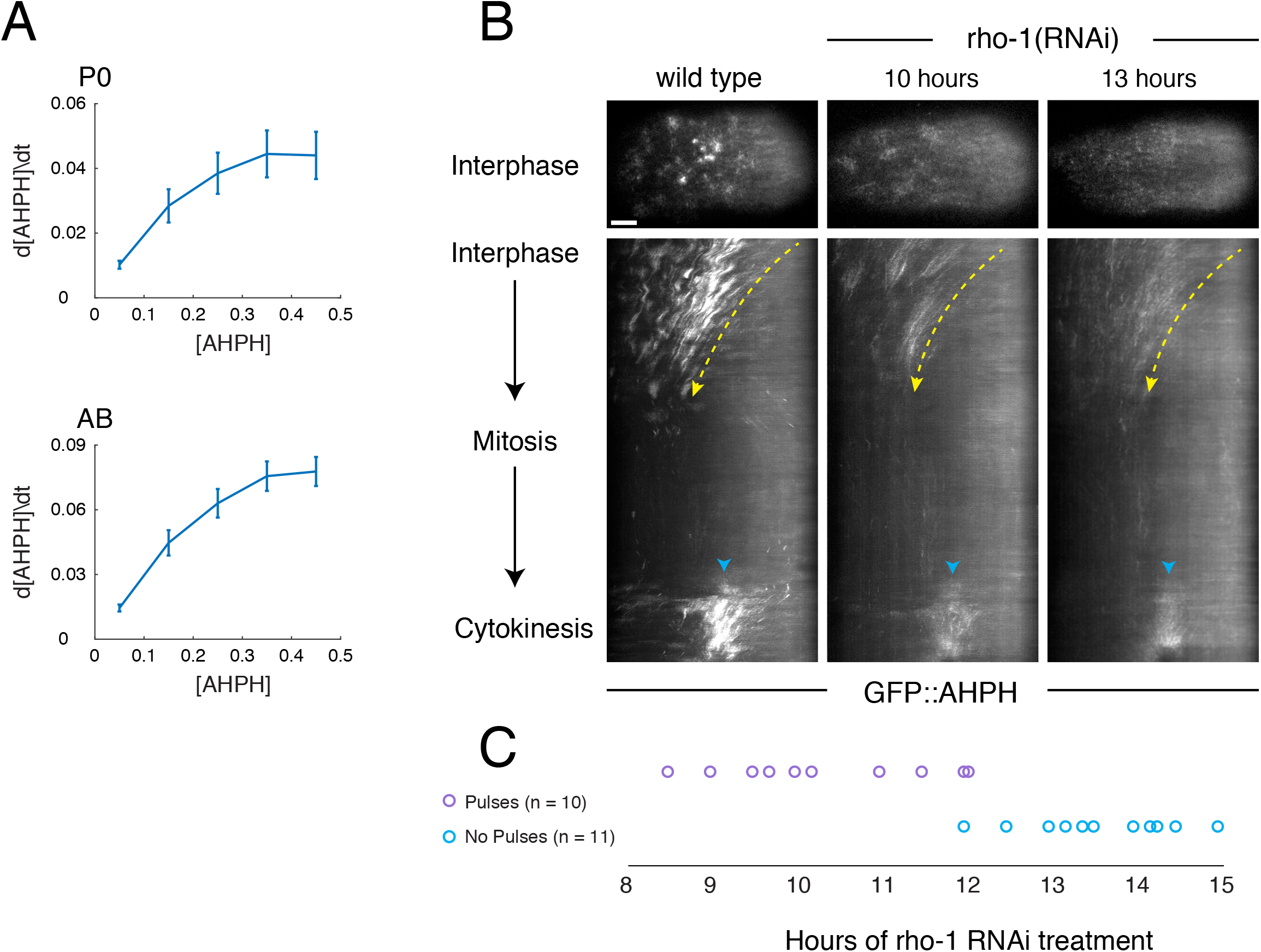
RhoA activation is autocatalytic. (A) The time derivative of normalized RhoA activity (GFP::AHPH) plotted vs. normalized activity during the early phase of pulse initiation in P0 (top panel, n = 40 pulses) and AB (bottom panel, n = 41 pulses). Error bars: 95% confidence interval. (B-C) Analysis of pulse dynamics in embryos progressively depleted of RHO-1 by RNAi. (B) Top panels show GFP::AHPH distributions in interphase embryos from mothers subjected to no (wild type), 10 hours and 13 hours of *rho-1(RNAi)*. Middle panels show kymographs from the same embryos illustrating spatiotemporal dynamics of GFP::AHPH from interphase through cytokinesis. Dashed yellow lines indicate approximate pattern of cortical flow. Cyan arrowheads indicate accumulation of GFP::AHPH just prior to cytokinesis. (C) Timeline indicating the presence (magenta circles) or absence (cyan circles) of pulsing in embryos treated with *rho-1(RNAi)* for the indicated times, revealing an abrupt transition from pulsing to no pulsing at ~12 hours post-treatment. Scale bars = 5µm in B.

### Delayed accumulation of the Rho GAPs RGA-3/4 underlies pulse termination

What terminates RhoA activity during each pulse? Our results imply that local termination of RhoA activity at the end of a pulse does not require Myosin II activity or (by extension) its local inhibition by Myosin phosphatase (Piekny and Mains, 2002; Piekny et al., 2003; Diogon et al., 2007), nor is it timed by cortical disassembly. Previous studies identified the redundant RhoA GAPs RGA-3 and RGA-4 as inhibitors of RhoA activity during polarization and cytokinesis (Schonegg et al., 2007; Zanin et al., 2013; Schmutz et al., 2007; Tse et al., 2012). A YFP-tagged RGA-3 transgene accumulates at the cortex in early embryos (Schonegg et al., 2007), and simultaneous depletion of RGA-3 and RGA-4 leads to hyper activation of RhoA and hypercontractility during zygotic polarization (Schonegg et al., 2007; Schmutz et al., 2007; Tse et al., 2012). We wondered, therefore, if RGA-3/4 could provide delayed negative feedback to terminate RhoA activity during individual pulses.

To test this possibility, we first imaged embryos co-expressing GFP::RGA-3 and NMY-2::mKATE. Focusing on AB, and using 2-color analysis as above, we confirmed that GFP::RGA-3 is present throughout the cortex, but accumulates locally during individual pulsed contractions (Figure 8A-C, Movie S8). Significantly, GFP::RGA-3 and NMY-2::mKATE accumulated with very similar timing (Figure 8B). Using NMY-2::mKATE and NMY-2::RFP as common signals to co-align data for GFP::AHPH and GFP::RGA-3, we inferred that GFP::RGA-3 accumulates with a ~6 sec delay relative to GFP::AHPH, the rate of RhoA activation peaks before the onset of GFP:RGA-3 accumulation, and rapid accumulation of GFP::RGA-3 coincides with deceleration and then reversal of RhoA activation (Figure 8C, bottom). Together, these observations suggest that delayed accumulation of RGA-3/4 plays a key role in terminating each pulse of RhoA activity. To test this further, we created a strain in which GFP::AHPH and NMY-2::mKATE were co-expressed in *rga-3;rga-4* (hereafter *rga-3/4*) double mutant embryos (Zanin et al., 2013). Consistent with previous reports (Schonegg et al., 2007; Zanin et al., 2013; Schmutz et al., 2007; Tse et al., 2012), during polarity establishment in *rga-3/4* double mutant zygotes, we observed hyper-accumulation of GFP::AHPH and hypercontractility, characterized by convulsive contractions of the anterior cortex and rapid anterior directed cortical flows (Figure 8D, 2nd column, Movie S9). However, we could no longer detect local pulses of GFP::AHPH in these embryos. To exclude the possibility that rapid flows sequester factors required for pulsed contractility to the extreme anterior pole, we used partial depletion of the myosin regulatory light chain (MLC-4) to attenuate contractility and cortical flows in *rga-3/4* double mutant zygotes or in control zygotes that were doubly heterozygous for *rga-3* and *rga-4* (Figure 8D, Movie S9). In *rga-3/4* heterozygotes partially depleted of MLC-4, cortical flows were sharply reduced, but pulsed accumulation of GFP::AHPH could be readily detected (Figure 8D, 3rd column, Movie S9). By contrast, in *rga-3/4* double mutant zygotes partially depleted of MLC-4, cortical flows during polarity establishment phase were slower than observed in wild type embryos; GFP::AHPH was uniformly enriched, but we did not observe local pulses of GFP::AHPH accumulation (Figure 8D, 4th column, Movie S9). Together, these data suggest that negative feedback through delayed accumulation of RGA-3/4 plays a key role in terminating local pulses of RhoA activity.

**Figure 8.**
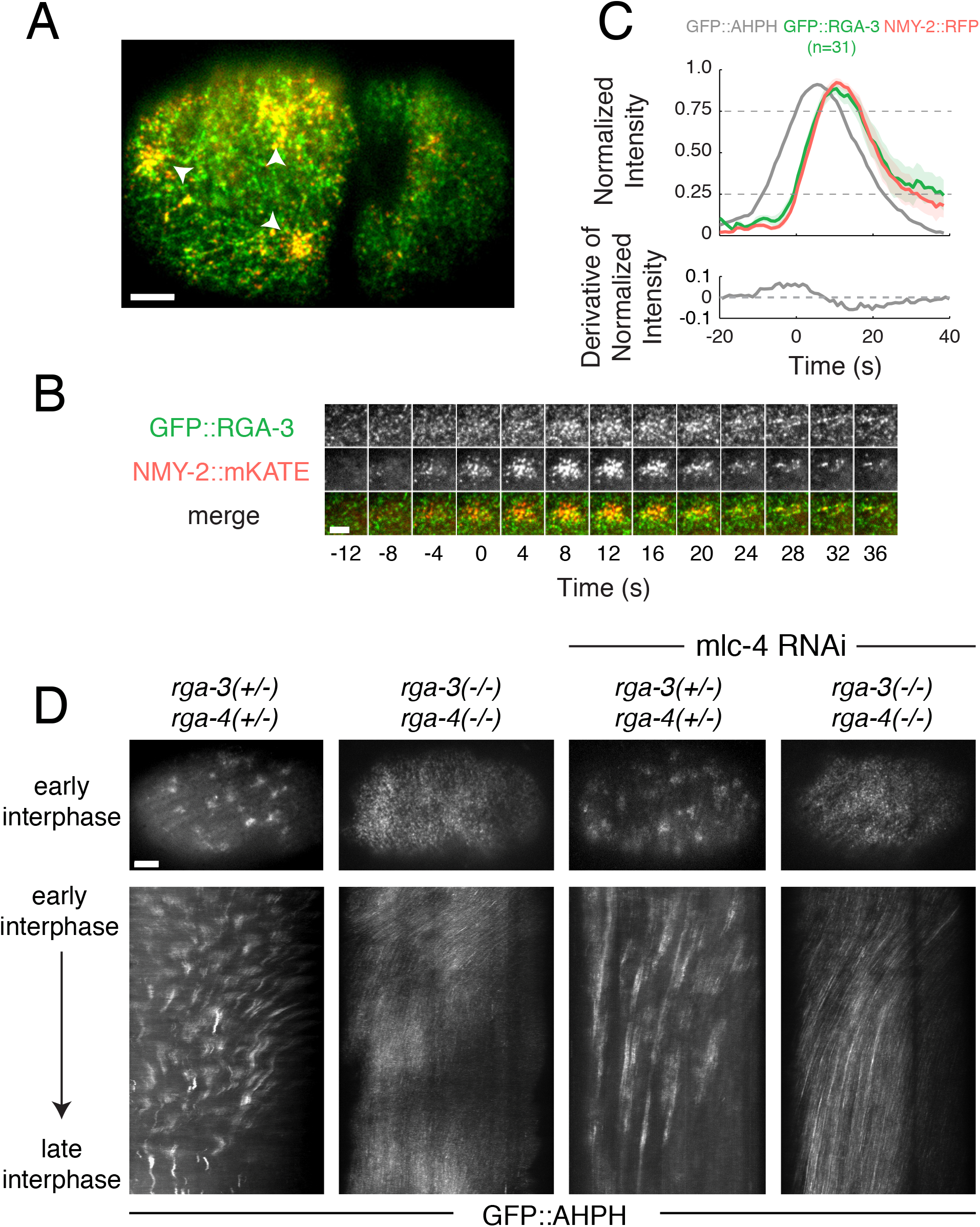
Delayed accumulation of RGA-3/4 mediates negative feedback required for pulse termination. (A) Micrograph of a 2-cell stage embryo expressing GFP::RGA-3 (green) and NMY-2::RFP (red). (B) Temporal dynamics of a single pulse. (C) Top: Averaged normalized fluorescence intensities vs. time for NMY-2::RFP and GFP::RGA-3 from two-color data, co-aligned with respect to the time at which NMY-2::RFP reaches 25% threshold. The averaged normalized fluorescence intensity of GFP::AHPH, co-aligned with NMY-2::RFP. Bottom: The averaged time derivative of the normalized GFP::AHPH intensity, again co-aligned using NMY-2::RFP. Hued regions report 95% confidence intervals. (D) Top: distributions of GFP::AHPH during interphase in zygotes with the indicated genotypes. Bottom: kymographs showing patterns of GFP::AHPH distribution and redistribution during interphase for the same genotypes. Scale bars = 5µm in A,D; 3µm in B.

### RGA-3/4 colocalizes with F-actin and requires F-actin for cortical association

Recent work suggests that F-actin accumulation inhibits RhoA activity to promote excitable dynamics in echinoderm and frog oocytes and embryos (Bement et al., 2015). We wondered if F-actin might play a similar role in *C. elegans* embryos by promoting recruitment of RGA-3/4. Consistent with this possibility, two-color live imaging of embryos co-expressing GFP::RGA-3 and mCherry::Lifeact (a marker for F-actin (Pohl et al., 2012)) revealed extensive overlap of GFP::RGA-3 and mCherry::Lifeact signals throughout the cortex (Figure 9A,D), and during individual pulses (Figure 9B,E), in both AB and P0 cells. Scatterplots showing pixelwise comparison of the two signals (Figure 9C,F), and measurements of Pearson’s (r) and Mander’s (M_RGA_) correlation coefficients over the entire cortex (P0: r = 0.9, M_RGA_ = 0.98; AB: r = 0.8; M_RGA_ = 0.96), or during individual pulses (P0: r = 0.84 +/-0.03, M_RGA_ = 0.97+/-0.02; AB: r = 0.65 +/-0.05; M_RGA_ = 0.89 +/-0.03; mean +/-SD; n = 10 pulses), confirmed strong colocalization.

**Figure 9.**
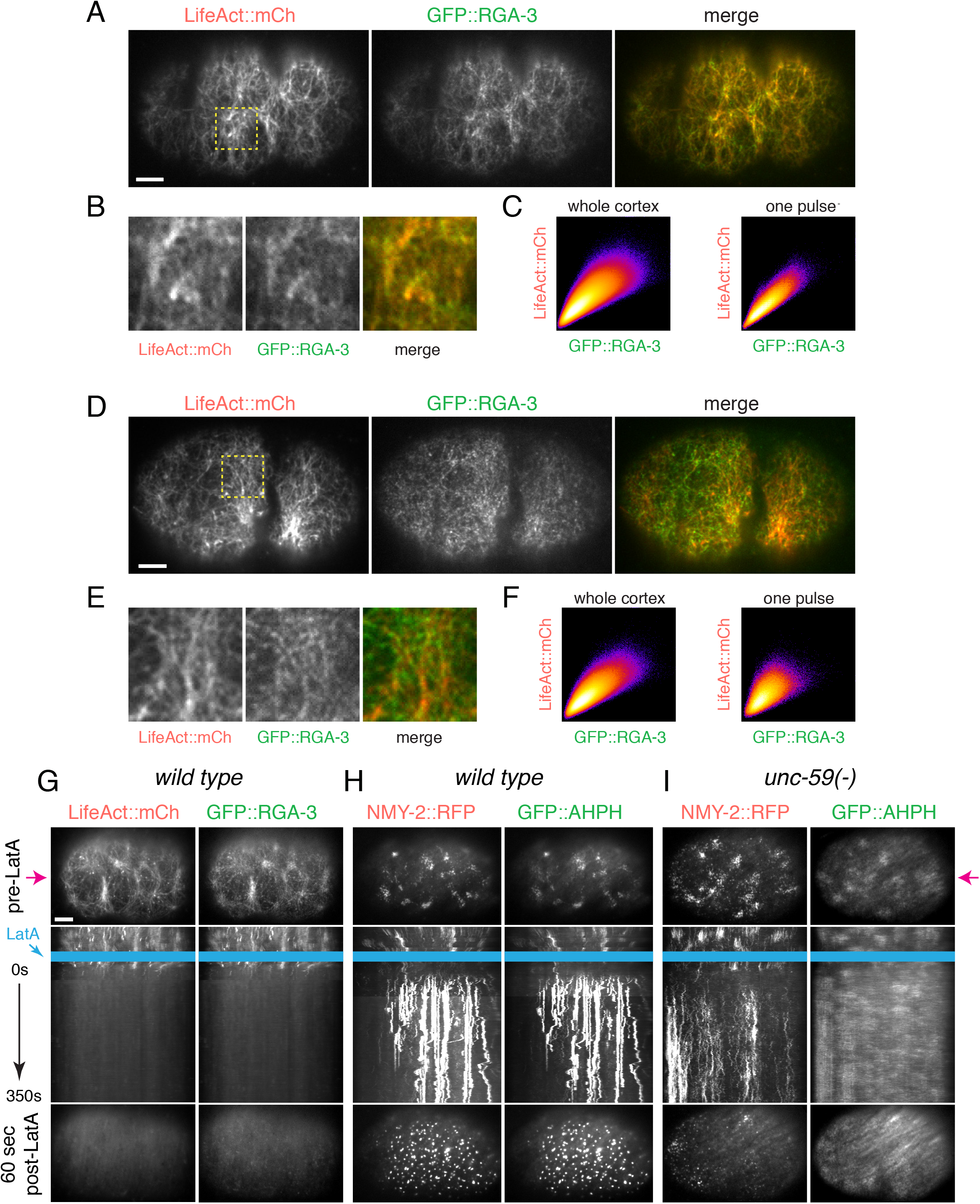
Cortical RGA-3/4 localization depends on F-actin. (A) Micrographs of a zygote co-expressing GFP::RGA-3 and mCherry::LifeAct. (B) Magnified view of the pulsing region indicated by the dashed yellow box in (A). (C) Scatter plots showing pixel-wise comparisons of GFP::RGA-3 and mCherry::LifeAct intensities for (left) the entire cortex of the zygote shown in A over 50 consecutive frames, or (right) for 50 consecutive frames in the region of cortex undergoing a pulse shown in B. (D) Micrographs of a two-cell embryo co-expressing GFP::RGA-3 and mCherry::LifeAct. (E) Magnified view of the pulsing region indicated by the dashed yellow box in (D). (F) Scatter plots showing pixel-wise comparisons of GFP::RGA-3 and mCherry::LifeAct intensities for (left) the entire AB cortex over 50 consecutive frames or (right) for 50 consecutive frames in the region of cortex undergoing a pulse shown in E. (G) Wild type zygote co-expressing GFP::AHPH and mRFP::LifeAct before (top) and ~90s after (bottom) treatment with 10µM Latrunculin A. Middle panel shows a kymograph taken from the region indicated by magenta arrows. Blue rectangle indicates approximate time of application of Latrunculin A. (H) Wild type zygote co-expressing GFP::AHPH and mRFP::Myosin II before (top) and ~90s after (bottom) treatment with 10µM Latrunculin A. Kymograph format as in G. (I) Homozygous *unc-59* (e261) mutant zygote co-expressing GFP::AHPH and NMY-2::RFP before (top) and ~90s after (bottom) treatment with 10µM Latrunculin A. Kymograph format as in G. Scale bars = 5µm in A,D,G.

To test whether F-actin is required for cortical localization of GFP::RGA-3, we treated permeabilized zygotes (Carvalho et al., 2011; Olson et al., 2012) co-expressing mCherry::Lifeact and GFP::RGA-3 with Latrunculin A (LatA) to rapidly depolymerize F-actin. Acute treatment with 10 µM LatA lead to a rapid collapse of the cortical F-actin network and complete disappearance of F-actin from the cortex within ~60 seconds. GFP::RGA-3 disappeared from the cortex with nearly identical timing, and remained closely associated with the F-actin network as it collapsed (Figure 9G, Movie S10, n = 6 embryos). These data suggest that cortical localization of RGA-3 depends on continuous physical association with F-actin. In contrast, upon acute depolymerization of F-actin, RFP::NMY-2, GFP-AHPH, GFP::Anillin, and GFP::Septin became rapidly enriched within dense punctate structures (Figure 9H; Figure S5A,B; data not shown), similar to what has previously been described in other contexts (Hickson and O’Farrell, 2008), suggesting that none of these other downstream targets of RhoA are sufficient for cortical recruitment of RGA-3 (at least in the absence of an intact cortical actin network).

We could no longer detect pulsed accumulation of AHPH after acute treatment of permeabilized zygotes with LatA. However this could be due to the sequestration of RhoA itself, or the biosensor, into dense clusters, as described above. We found that GFP::AHPH is no longer enriched in dense clusters after LatA treatment in embryos mutant for the Septin *unc-59*, (Figure 9I). Pulsed accumulation of GFP::AHPH and NMY-2::RFP persisted in *unc-59* mutant zygotes (Figure S5D,E). However, acute treatment of permeabilized *unc-59* zygotes with LatA lead to rapid accumulation of a diffuse pool of GFP::AHPH throughout the cortex and a loss of pulsing (Figure 9I), analogous to what we observe in *rga-3/4* double mutant zygotes.

### Fast positive and delayed negative feedback involving RhoA and RGA-3/4 can account quantitatively for locally pulsatile RhoA dynamics

Our data suggest that in the early *C. elegans* embryo, locally excitable RhoA dynamics arise independently of myosin activity through a combination of fast positive feedback on RhoA activity and delayed negative feedback via local F-actin-dependent recruitment of RGA-3/4 (Figure 10A). To test this idea, we asked if a simple model based on these feedbacks with parameters constrained by experimental measurements, predicts excitable dynamics. (Figure S7, materials and methods). We formulated the model as a pair of ordinary differential equations, describing local rates of change in active RhoA and RGA-3/4, based on the following assumptions: (a) RhoA is activated at a basal rate, and feeds back positively to promote further RhoA activation, (b) RhoA feeds forward through F-actin assembly to promote local, reversible, association of RGA-3/4 with the cortex and (c) RGA-3/4 acts as a GAP to promote local inactivation of RhoA. Consistent with our experimental observations (Figure 6A, Figure S7A), we assumed that autoactivation of RhoA is a saturating function of RhoA activity, represented by a Hill function with Hill coefficient n = 1. We assumed that inactivation of RhoA by RGA-3/4 obeys Michaelis-Menten kinetics. To account for the observed delay between an increase in RhoA and the sharp onset of RGA-3/4 accumulation (Figure 7C), we assumed ultrasensitive dependence of RGA-3/4 accumulation rate on RhoA, with the steepness of the response governed by an exponent m (see Materials and Methods for mathematical details).

**Figure 10.**
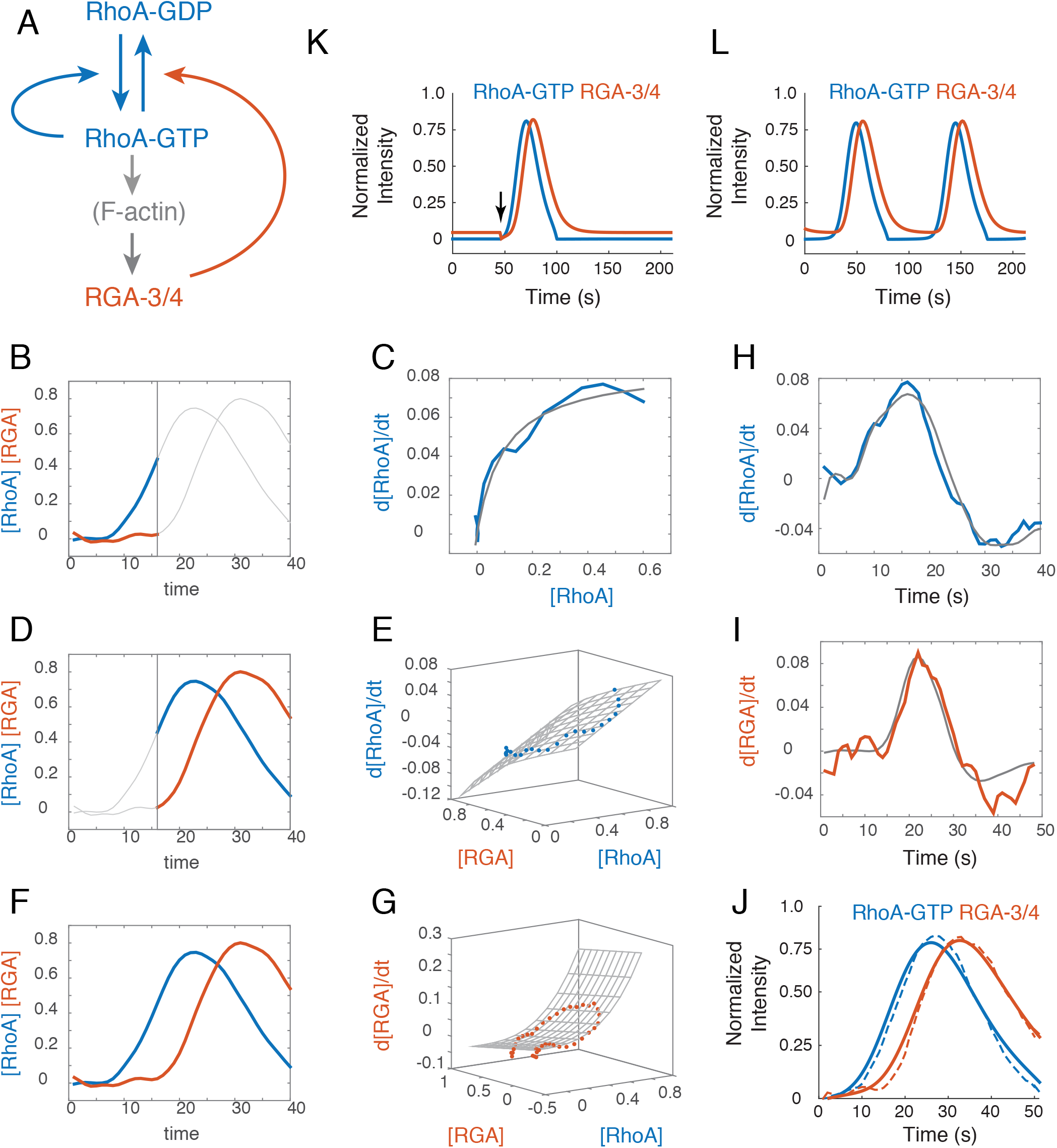
Autocatalytic RhoA activation and delayed negative feedback through RGA-3/4 is sufficient to produce locally excitable RhoA dynamics. (A) Schematic representation of a simple mathematical model for RhoA pulse dynamics, based on our results. (B) Comparison between measured (dashed lines) and simulated (solid lines) pulse dynamics for the case in which n = 1, 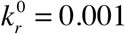, 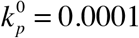, *K_GAP_* = 0.001. The remaining model parameters were estimated by fitting model equations to experimental data (*m* = 3.02, 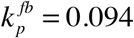, 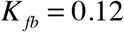, 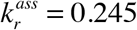, 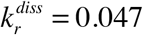); see Materials and Methods for details). (C) Simulated dynamics for the parameter values in (B) are excitable, with a stable rest state can be destabilized by a transient reduction of RGA-3/4 (vertical black arrow) to trigger a single pulse of RhoA activity. (D) A small change in the basal RGA-3 association rate from 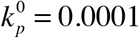 to 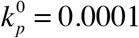 results in oscillatory dynamics, with pulses occurring at regular intervals

We set values for basal RhoA activation and RGA-3./4 recruitment rates to very low values, based on the slow rates of increase in GFP::AHPH and GFP::RGA-3 observed before the sharp upswing of each pulse (Figure 7C). Then we estimated values for the model’s remaining parameters by fitting the right hand sides of equations (1) and (2) to the relationships between dRhoA/dt and dRGA-3/4/dt and RhoA and RGA-3/4, inferred from co-aligned, averaged and normalized GFP::AHPH and GFP::RGA-3 intensities (Figure S7A-F; see Materials and Methods for details).

Given these parameter estimates, and depending on the exact values chosen for the basal RhoA activation and RGA-3/4 association rates, we observed one of two closely related behaviors: Excitable dynamics, characterized by a stable rest state (low RhoA and low RGA), from which the system can be induced to generate a transient pulse by a small decrease in RGA (Figure 9B) or a small increase in RhoA (not shown), and oscillatory dynamics, in which the stable rest state is lost and the system undergoes transient pulses at regular intervals (Figure 9C). The transition from excitable to oscillatory dynamics could be induced by either a small increase in basal RhoA activation or a small decrease in basal RGA activation. This is consistent with our observations in *nmy-2(RNAi)* embryos that some patches of cortex are quiescent while others exhibit repeated pulses of activity at regular intervals (yellow arrows in Figure 5A), and suggests that, in the absence of contractilty, the *C. elegans* cortex may be poised close to a transition between excitable (with a low threshold for excitation) and oscillatory dynamics. We conclude that a simple combination of positive and negative feedback loops, coupling local RhoA activity and RGA-3/4 accumulation, is in principle sufficient to explain pulsatile RhoA dynamics in early *C. elegans* embryos, independent of actomyosin contractility.

## Discussion

Pulsed contractility is a widespread mode of actomyosin contractility, but its mechanistic basis has remained poorly understood (Levayer and Lecuit, 2012; Gorfinkiel, 2016). Current models for pulsed contractility invoke mechanochemical feedback in which contractile forces produced by Myosin II couple in different ways with actomyosin assembly/disassembly to drive excitable or oscillatory dynamics. Proposed feedback mechanisms include tension-dependent motor binding kinetics (Ren et al., 2009; Effler et al., 2006; He et al., 2010; Luo et al., 2012), tension-dependent filament assembly/stabilization (Hayakawa et al., 2011; La Cruz and Gardel, 2015) or disassembly (Machado et al., 2014), tension-dependent activation of Myosin II via e.g. Ca++ (Kapustina et al., 2008) or RhoA (Koride et al., 2014), or modes of feedback in which local contraction advects and concentrates actomyosin and/or its upstream activators (Bois et al., 2011; Kumar et al., 2014; Munjal et al., 2015; Nishikawa et al., 2017). Here, we have identified a mechanism for pulse generation in early *C. elegans* embryos that does not require force production or redistribution of cortical factors by Myosin II. Using single molecule imaging and particle tracking analysis, we have shown that the rapid initial accumulation of F-actin and Myosin II begins well before the onset of contraction, at a time when the cortex is locally expanding; Redistribution of actomyosin by local contraction makes a minor contribution to the overall accumulation of actomyosin during each pulse. Instead, our data show that pulsed accumulation and disappearance of F-actin and Myosin II are determined primarily by local modulation of their assembly/recruitment and disassembly. Using two-color imaging, we have shown that during each pulse, a biosensor for active RhoA begins to accumulate well before RhoA’s downstream targets F-actin, Myosin II and Anillin. Active RhoA nearly reaches its peak level before the onset of contraction (Figure 4D), and then it begins to disappear well before its downstream targets. Significantly, locally pulsed activation of RhoA continues to occur on patches of cortex that contain only a few (1-2) particles of Myosin II, too few to produce significant local contractile stress. Thus a Myosin-independent RhoA pulse-generator underlies pulsed contractility in early *C. elegans* embryos.

One important caveat is that the ANI-1 AHPH domain may bind septin, or lipids, or other cortical targets in addition to active RhoA (Piekny and Maddox, 2010). Indeed, we observe what appears to be two phases of AHPH accumulation – a diffuse phase that accumulates early, and a second phase that accumulates later and colocalizes with NMY-2. The diffuse pool is likely to reflect RhoA-GTP that is available to bind RhoA affectors. Whether the myosin-colocalized pool bids active RhoA or other cortical factors, remains unclear. Importantly, the diffuse pool has very similar kinetics in wild type AB, and in polarization-deficient or myosin-depleted P0 cells, supporting the conclusion that a common myosin-independent mechanism for pulsed activation of RhoA that operates in both P0 and AB cells.

The pulses of RhoA activity described here and in other contexts (Munjal et al., 2015; Bement et al., 2015; Mason et al., 2016) are strikingly reminiscent of excitable behaviors found in other systems, such as action potentials in neuronal (Izhikevich, 2007) or cardiac cells (Luo and Rudy, 1991), or transient pulses of intracellular calcium release (Goldbeter, 1996), or pulses of actin assembly observed in motile cells (Weiner et al., 2007). Theoretical studies highlight two key ingredients for excitable dynamics: fast positive feedback and delayed negative feedback (Strogatz, 1994). The sharp acceleration of active RhoA accumulation that we observe during the early rising phase of individual pulses is a dynamical signature of fast positive feedback in which RhoA promotes its own activity. Stronger evidence that RhoA participates in a positive feedback loop that is essential for pulsing comes from our observation that depletion of RhoA below a certain threshold leads to an abrupt loss of pulsed contractions, while having minimal effects on other RhoA-dependent functions such as cortical flow during polarization (Motegi and Sugimoto, 2006; Schonegg and Hyman, 2006) or cytokinesis (Loria et al., 2012).

The mechanism for this feedback remains unclear. In AB cells, the rapid acceleration of RhoA activation during pulse initiation occurs before any visible accumulation of Myosin II, Anillin or F-actin, and in P0, depletion of NMY-2 or ANI-1 has little affect on the rising phase of RhoA activation. Thus, with the possible exception of F-actin in P0, it seems very unlikely that accumulation of these downstream targets makes a significant contribution to positive feedback. A more likely possibility is that RhoA feeds back through one or more of its upstream activators, such as ECT-2, CYK-4 and NOP-1 (Tse et al., 2012). In the zygote, pulsing is sensitive to depletion of ECT-2 and NOP-1 (Tse et al., 2012; Naganathan et al., 2014). During cytokinesis, active RhoA can act as a cofactor to promote trans-activation of the RhoGEF ECT-2 by the RhoGAP CYK-4 (Zhang and Glotzer, 2015). While CYK-4 is not required for pulsed activation of RhoA during polarization, it is possible that RhoA could feedback through NOP-1, a protein of unknown activity that is required for RhoA activation during interphase in P0 and AB (Tse et al., 2012). Identifying the molecular mechanism(s) for this feedback is an important goal for future studies.

Regarding delayed negative feedback, direct comparison of our single molecule and two-color imaging data (compare Figures 3 and 4) argues strongly against a mechanism in which pulses are terminated by actomyosin disassembly. Both total and diffuse GFP::AHPH signals peak and begin to fall sharply at a time when F-actin and Myosin II disassembly rates are lowest (compare Figures 3&4). Indeed, our data suggest that local accumulation of ANI-1 downstream of RhoA stabilizes Myosin II (and possibly also F-actin) to prolong the contraction phase of each pulse. Instead, our data suggest that the redundantly acting RhoGAPs RGA-3/4 play a key role in providing the delayed negative feedback that terminates RhoA pulses. RGA-3/4 act as GAPs towards RHO-1 in vitro (Schonegg et al., 2007), and loss of RGA-3/4 leads to hyperactivation of RhoA in vivo (Tse et al., 2012). We find that during each pulse, RGA-3 accumulates with a delay of ~6 seconds relative to active RhoA. Significantly, ***the rate*** of active RhoA accumulation peaks, and begins to fall, just as GFP::RGA-3 begins to accumulate, suggesting that rapid accumulation of RGA-3/4 plays a key role in timing the end of each RhoA pulse. Consistent with this possibility, RhoA activity is uniformly high, and pulsatile RhoA activation is completely abolished, in *rga-3/4* double mutant zygotes, even when contractility is attenuated to prevent sequestration of RhoA activators by hyperactive cortical flow. Whether factors other than RGA-3/4 contribute to terminating RhoA pulses remains unclear.

Together, these data suggest a model for locally excitable RhoA dynamics in which RhoA feeds back positively to promote its own activation, and feeds back negatively with a delay through RGA-3/4 to promote its own inactivation. Indeed, when we formulate this model mathematically, and constrain model parameter values to match the local dependencies of RhoA and RGA-3/4 accumulation rates on levels of RhoA and RGA-3/4 inferred from two-color imaging data, the model predicts locally pulsatile RhoA dynamics, and small tunings of the model’s parameters mediate interconversion between excitable dynamics and spontaneous oscillations. This simple modeling exercise establishes an internally consistent hypothesis for pulsatile contractility that can be confirmed and extended by future experiments.

What governs the recruitment of RGA-3/4 during each pulse? We have found that GFP::RGA-3 co-localizes broadly and extensively with cortical F-actin in both P0 and AB. RGA-3/4 accumulates with the same timing as F-actin during each pulse, and depolymerizating F-actin abolishes this accumulation. This suggests a specific mechanism for delayed recruitment of RGA-3/4 in which RhoA promotes increased local F-actin assembly (potentially through the formin Cyk-1 (Severson et al., 2002)), and F-actin in turn recruits RGA-3/4. Interestingly, a recent study (Bement et al., 2015) suggests that RhoA and cortical F-actin form an excitable circuit, with RhoA as activator and F-actin as inhibitor, that propagates cortical waves of RhoA activity and F-actin assembly in oocytes and embryonic cells of frogs and echinoderms. The mechanism(s) by which F-actin feeds back to inactivate RhoA in these cells remains unknown, but our observations in *C. elegans* support to the idea, proposed by Bement et al, 2015, that a RhoGAP homologous (or analogous) to RGA-3/4 may be recruited by F-actin to mediate negative feedback in frog and echinoderm cells. A similar mechanism has recently been reported in the context of secreting in Drosophila larval salivary glands (Segal et al., 2018), and a similar circuit design may underlie the propagation of actin waves observed in many motile cells (reviewed in (Allard and Mogilner, 2013)).

It is also interesting to compare our observations to those made recently in the Drosophila germband (Munjal et al., 2015). In germband cells, pulsed accumulation of a biosensor (GFP fused to the AHPH domain of Drosophila anillin) appears to coincide with the local accumulation of F-actin and Myosin II, and with the onset of contraction, and it is abolished by inhibition of Rho Kinase, an upstream activator of Myosin II. In *C. elegans*, by contrast, pulsed accumulation of GFP:AHPH precedes actomyosin accumulation and the onset of contraction by many seconds and persist in the almost complete absence of Myosin II.

To some extent, these differences could reflect the imaging methods used to detect the RhoA biosensor. Using near-TIRF imaging in *C. elegans* embryos, we detect two pools of the biosensor: a diffuse pool that begins to accumulate well before Myosin II, and a second more punctate pool whose distribution strongly overlaps with Myosin II (Figure 4B, Figure S3F&G, Figure S5. Based on studies in other cells (Weiner et al., 2007), the diffuse pool may be more difficult to detect using confocal microscopy. Thus it remains possible that a diffuse pool of active RhoA accumulates before actomyosin in the Drosophila germband, but escapes detection by confocal microscopy.

An alternative idea is that pulsed contractility is governed by locally excitable RhoA dynamics in both systems, but that different forms of positive feedback may contribute differently to driving the rapid upswing of RhoA activity, and that different mechanisms may operate to trigger pulses (by driving RhoA activity above a threshold for excitation). For example, in the Drosophila germband, a mode of feedback in which local contraction advects and concentrates active RhoA (or upstream activators) may be required to initiate pulses, whereas in *C. elegans*, local fluctuations in RhoA or RGA-3/4 levels may be sufficient to do so in the absence of contractility. Importantly, in the Drosophila germband, as in *C. elegans*, advection/contraction coupling accounts for only a fraction of the total accumulation of active RhoA during each pulse; thus other modes of positive feedback must also make a significant contribution.

More generally, we hypothesize that the different modes of RhoA excitability that have been described in frog, echinoderm, *C. elegans* and *Drosophila* embryos share a deeper underlying mechanistic origin. We suggest that a comparative analysis of mechanisms for pulsing in these and other systems will be a very fruitful way to uncover core conserved circuitry for pulse generation and to understand the ways in which this core circuitry is tuned or accessorized in different contexts to achieve different functional outcomes.

## Materials and Methods

### *C. elegans* culture and strains

We cultured *C. elegans* strains at 22˙C under standard conditions (Brenner, 1974) Table S1 lists the mutations and transgenes used in this study. Unless otherwise specified, strains were provided by the Caenorhabditis Genetics Center, which is funded by the National Institutes of Health (NIH) National Center for Research Resources.

### RNA interference

RNAi was performed by the feeding method as previously described (Timmons et al., 2001). Bacteria targeting *nmy-2, spd-5, rho-1, perm-1, arx-2, ani-1* and *mlc-4* were obtained from the Kamath feeding library (Kamath et al., 2003). The L4417 plasmid targeting the entire GFP sequence (generated by the Fire lab and available at (Rauzi et al., 2010; Levayer and Lecuit, 2013; He et al., 2010)) was transformed into HT115(DE3) bacteria. For the Myosin depletion experiments, L4 larvae co-expressing GFP::AHPH and NMY-2::mKate2 were transferred to *nmy-2* RNAi feeding plates 24-30 hours before imaging. Strong depletion of myosin was verified by strong loss of cortical NMY-2::mKate2. For the ANI-1 depletion experiments, L4 larvae co-expressing GFP::AHPH and NMY-2::RFP, were transferred to *ani-1* RNAi feeding plates 30-36 hours before imaging. For experiments involving *spd-5* RNAi, L4 larvae were transferred to feeding plates for 24-30 hours before imaging. For the RhoA depletion experiments, synchronized young adults were transferred to *rho-1* RNAi plates 8-16 hours before imaging. For experiments involving *mlc-* 4 RNAi, synchronized young adults were transferred to feeding plates for 12-16 hours before imaging. For the latrunculin A experiments, late L4 larvae were transferred to *perm-1* RNAi plates 16-24 hours before imaging. For experiments involving *arx-2* RNAi, L4 larvae were transferred to feeding plates for 30-36 hours before imaging.

### Microscopy

We mounted embryos as described previously (Robin et al., 2014) on glass slides under #1.5 coverslips in 3-5µl of standard Egg Salts containing ~100 uniformly sized polystyrene beads (18.7 ± 0.03 μm diameter, Bangs labs, #NT29N). The beads acted as spacers and allowed us to achieve uniform compression of the embryo surface across experiments (Robin et al., 2014).

We performed all imaging on a Nikon ECLIPSE-Ti inverted microscope equipped with a Ti-ND6-PFS Perfect Focus Unit. A laser merge module (Spectral Applied Research) controlled fast, tunable delivery of 481nm and 561 nm laser excitation from 50mW solid state lasers (Coherent Technology) to a motorized TIRF illuminator. We adjusted laser illumination angle to achieve near-TIRF illumination (Tokunaga et al., 2008). We collected images using a Nikon CFI Apo 1.45 NA oil immersion TIRF objective combined with 1.5 intermediate magnification onto an Andor iXon3 897 EMCCD camera. All image acquisition was controlled using Metamorph software.

### Single-molecule imaging

We performed single molecule imaging as described previously (Robin et al., 2014). For NMY-2::GFP, we used a combination of RNAi against GFP and mild photobleaching in wide field illumination mode to reduce surface densities of GFP-tagged transgenic proteins to single molecule levels. For GFP::Actin, which is expressed at very low levels in the strain that we used, we used mild photobleaching alone. For GFP::Actin and NMY-2::GFP, we imaged single molecules using 10% laser power (~0.16 μW.μm^−2^), with 100ms exposures in continuous streaming mode (GFP::Actin and NMY-2::GFP), yielding an approximate photobleaching rate of ~0.05 s^−1^ (Robin et al., 2014).

### Analysis of F-actin and Myosin II turnover

In previous work, we compared two methods for estimating local F-actin disassembly rates from single molecule data (Robin et al., 2014). The first method (smPReSS) estimates average disassembly rates in a local region by fitting kinetic models to the approximately exponential decay in particle densities measured during photobleaching, assuming steady state conditions. The second method relies on single molecule detection and tracking and infers appearance and disappearance events directly from single molecule trajectories. We showed that under steady state conditions, and when Myosin II is inhibited to remove the effects of local contraction and cortical flow, these two methods yield estimates of local disassembly that agree to within 20%. During pulsed contractions, the steady state assumption is not valid and the effects of cortical flow cannot be ignored. Therefore, in this work, we relied exclusively on the particle tracking method to measure local appearance, disappearance and motion of single molecules.

In preliminary analyses, we found that single molecules of GFP::Actin and NMY-2::GFP move sufficiently slowly during pulsed contractions that we could obtain marginally better results by pre-averaging ten consecutive frames of raw data to produce sequences of images at one-second intervals. We performed single molecule detection and tracking on this pre-averaged data using a Matlab implementation (http://people.umass.edu/kilfoil/downloads.html) of the Crocker-Grier method (Crocker, 1996; Pelletier et al., 2009). We then inferred single molecule appearance and disappearance events and frame-to-frame single molecule displacements directly from the single molecule trajectories.

### Measuring local deformations from single molecule data

The key goal of our single molecule analysis was to distinguish the relative contributions of local assembly/disassembly and local deformation/flow to changes in local density during pulsed contractions. To do so, it was essential to follow dynamic changes in assembly/disassembly on a moving and contracting patch of cortex, i.e. in a material (Lagrangian) frame of reference. We used single molecule displacements to identify and track regions of cortex undergoing pulsed contractions as follows:

### Tracking a polygonal region of interest (ROI) during individual pulses

For each pulse, we identified a reference time point at/near the onset of contraction by visual inspection of the time lapse sequence. At this time point, we identified manually an elliptical region containing the patch of cortex undergoing contraction (Figure S1A). We computed the smallest polygon (the “convex hull”) containing all the particles detected within the elliptical region (Figure S1B). Each vertex of the reference polygon was thus associated with a single molecule detected on the cell surface. To propagate the polygonal ROI forwards and backwards in time, we computed the frame-to-frame displacement of each of its vertices, either from the displacement of a vertex-associated molecule or (once the molecule disappears) from a weighted average of the frame-to-frame displacements of nearby particles (Figure S1C). We then measured local deformation and turnover within this polygonal ROI as follows:

### Measuring local deformation and turnover during individual pulses

We compared three different measures of local compression (or dilation) within the polygonal ROI from frame to frame (Figure S2A): the change in normalized surface area, a particle-based strain rate and a material strain rate. We computed the change in surface area s_A_ as the time-derivative of the normalized area of the polygonal ROI:

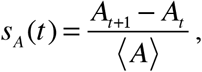

where ⟨*A*⟩ is the mean surface area taken over all frames in the pulse sequence. To compute a particle-based strain rate, for each particle in the polygonal ROI, we computed the average normalized change in distance between that particle and its near-neighbors:

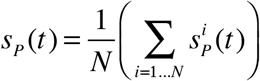

where 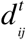 is the distance between particle i and a near neighbor particle j at time t and the sum is taken over all neighbor particles within a disk of radius 20 pixels centered on particle i. We then averaged over all particles in the ROI to obtain a particle-based strain rate for the entire ROI:

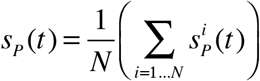

Finally, to compute a material strain rate s_M_, we used a linear least squares regression method to estimate the local gradient of particle velocities (Landau and Lifshitz, 1987). We then decomposed the resulting velocity gradient tensor into anti-symmetric (rotation) and symmetric (strain rate) components. We then took one half the trace of the symmetric strain rate tensor as a measure of local compressive strain. In practice, we found that all three methods yielded very similar results regarding the magnitude and timing of contractions (Figure S2A). We report results based on the particle-based strain rate in Figures 2 and 3.

### Measuring turnover

To quantify turnover rates for F-actin or Myosin II within the polygonal ROI, for each time point t, we measured the area of the ROI (A_t_), the number of particles N_t_, their density 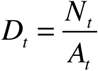, and the number of appearance and disappearance events that occurred within the ROI between time t and 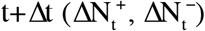. We quantified the mean appearance rate and the mean disappearance rates within the ROI as :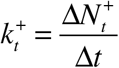 and 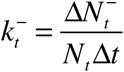. We computed the change in actin density within a polygonal ROI at time t as Δ*D_t_* = *D*_*t*+Δ*t*_ − *D_t_*. We estimated the contribution to the change in density from deformation of the ROI to be Δ*D*_*deformation*_ = −*D_t_s_P_*, where *s_P_* is the particle-based strain rate measured as described above, and the contribution from turnover (i.e. net imbalance of appearance and disappearance) to be 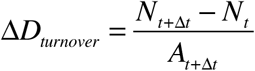, such that ∆D_t_ = ∆D_deformation_ + ∆D_turnover_.

### Two-color imaging, pulse tracking, and analysis

We performed two-color imaging using the imaging system described above with near-TIRF illumination. We performed the initial steps of image processing, pulse identification and extraction, using the software package ImageJ (http://imagej.nih.gov/ij/). For all subsequent steps, including pulse tracking, intensity measurements, data normalization and alignment across multiple pulses, we used custom functions written in MATLAB (http://www.mathworks.com).

We imaged embryos co-expressing GFP- and RFP-tagged transgenes by alternating 100 ms exposures with 488nm and 561nm excitation, thus giving 5 two-color frames per second. We used 25% maximum laser power ( ≈ 0.4 *µWµm* ^−2^) for each channel. For subsequent analysis, we averaged over five consecutive frames to obtain a single image for each channel at one-second intervals. We limited our analysis to individual pulses that moved very little during the period leading up to the onset of contraction. We used ImageJ to extract a subregion containing each pulse of interest. With the exception of Myosin-depleted embryos, we used NMY-2::RFP as a reference signal to track the location of the pulse through time. For the analysis of Myosin-depleted embryos, we used GFP::AHPH as the reference signal.

We used the reference signal to identify and track a moving region of interest associated with each pulse as follows (Figure S3A): First, we smoothed each frame of the image sequence using a gaussian filter, sigma = 2-3 μm). We then thresholded the smoothed image to identify regions of interest (ROIs) associated with the pulse in consecutive frames. We used the same value of sigma and the same threshold for all frames, and we chose these values such that each ROI in the sequence was simply connected and such that the largest ROI in the sequence was approximately the same size as the region of strong signal accumulation near the peak of the pulse in the unprocessed data, as viewed by eye.

To measure signal intensities vs. time during a pulse, we first used MATLAB to determine the centroid of each of the ROIs identified above to obtain a sequence of centroid positions *C_t_* = (*x*_*t*_, *y*_*t*_). We extended this sequence backwards in time using the first measured centroid position C_first_ and extended it forwards in time using the last measured centroid position C_last_. We then used a single reference ROI, centered on positions [*C_first−N_*,…, *C_last+N_*], to measure a sequence of GFP and RFP intensities over time. We compared two different reference ROIs: (i) the largest ROI measured in the sequence, which corresponds roughly to peak accumulation of Myosin II and (ii) a “bounding box”, defined to be the smallest square region containing this largest ROI. Because the results did not differ significantly, we used the bounding box ROI for all measurements reported here.

For each probe, we measured the mean signal intensity in the entire ROI over time. We then normalized these data for individual pulses using the equation 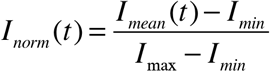, where *I_mean_*(*t*) is the mean intensity in the ROI at time t, *I_min_* is the minimum mean intensity in the ROI before the onset of contraction, *I*_max_ is the maximum mean intensity in the ROI measured over the entire sequence [*C_first−N_*,…, *C_last+N_*]. Finally, we aligned data across multiple pulses with respect to the time point at which NMY-2::RFP crossed 25% of its normalized maximum intensity (Figure S3C). The mean was calculated with a 95% confidence interval.

To partition the AHPH::GFP signal into “diffuse” and “myosin colocalized” fractions, we defined a “myosin colocalized” mask to be the collection of pixels within the ROI in which the raw NMY-2::RFP signal was greater than a threshold percentage (75% - 95%) of its peak intensity, and the “diffuse” mask to be its complement. (Figure S3D top). We used these masks to decompose the total GFP::AHPH signal into “diffuse” and “myosin-colocalized” pools (Figure S3D bottom). We then partitioned the total normalized intensity within the ROI into diffuse and myosin-colocalized signals given by 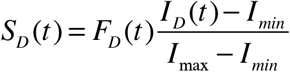 and 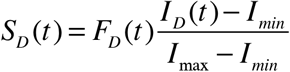, where F_<u>*D*</u>_(t) and F_*C*_(t) are the fraction of pixels, and I_<u>*D*</u>_(t) and I_C_(t) are the mean intensities, in the diffuse and myosin-colocalized regions respectively (Figure S3E,F). The timing with which the diffuse and myosin-colocalized signals rose and fell was insensitive to the exact threshold. We used 95% for the results reported here.

We performed a number of additional controls to assess the sensitivity of our results to variation across strains, or the details of pulse identification, tracking and intensity measurements. First, we compared the kinetics of Myosin II accumulation during pulses in embryos co-expressing NMY-2::GFP and NMY-2::RFP and confirmed that the dynamics of accumulation were essentially identical after normalizing for differences in expression level and probe brightness (Figure 4E,4F, Figure S3G,H). Second, we confirmed that the dynamics of Myosin::RFP accumulation was essentially identical across the different two-color strains that we used (Figure S3G,H). Finally, we verified that our measurements of the rate and relative timing of accumulation of different signals were largely insensitive to differences in the size of the box/blobs used (Figure S3, data not shown).

### Kymograph analysis

To produce the kymographs shown in Figures 4-8, Figure S4, and Figure S6, we aligned images so that the AP axis of the embryo coincided with the horizontal (x) image axis. We selected rectangular regions aligned with the x image axis, whose width (in x) coincided with the embryonic region of interest and whose height (in y) was 10-20 pixels. From the original image stack, we extracted an xyt substack corresponding to this rectangular region; we used ImageJ’s reslice tool to reslice this stack with respect to the xt plane, then we used a maximum intensity projection to collapse the individual slices in y to obtain a kymograph in x vs. t.

### Colocalization analysis

We analysed colocalization of mCherry::Lifeact and GFP::RGA-3 signals using the ImageJ plugin Coloc_2 (https://imagej.net/Coloc_2). Given a pair of image stacks representing the two signals for a sequence of frames within a region of interest, we first scaled the data for each stack to lie between 0 and 255, then used Coloc 2 to compute Pearson’s correlation coefficient and an associated significance value, and to estimate Mander’s correlation coefficients with automated threshold determination.(Costes et al., 2004)

### Modelling RhoA pulse generation

We built a simple ordinary differential equation model for RhoA pulse generation at a single point in space based on autocatalytic activation of RhoA and delayed negative feedback via RhoA-dependent recruitment of RGA-3/4. We started with the following assumptions:

1. RhoA is activated at a constant basal rate.
2. Active RhoA feeds back to promote further RhoA activation at a rate that can be described as a Hill function of RhoA density.
3. RGA-3 and RGA-4 can be treated as a single species (RGA-3/4) that acts as a GAP to promote local inactivation of RhoA.
4. Active RhoA promotes local F-actin assembly; RGA-3/4 binds F-actin from an abundant cytoplasmic pool, and dissociates from F-actin at a constant rate. Because RGA-3 and F-actin accumulate with very similar kinetics, we did not model F-actin directly. Instead, we assumed that RGA-3/4 binds the cortex at a constant basal rate plus a rate that depends on the local density of active RhoA, and that RGA-3/4 dissociates from the cortex at a constant rate.
5. To capture the observed delay between the sharp upswing in RhoA activity and the onset of F-actin and RGA-3/4 accumulation (Figure 4D, Figure 7B), we assumed ultrasensitive dependence of RGA-3/4 recruitment rate on RhoA activity.

With these assumptions, letting p represent the density of RhoA and r represent the density of RGA-3/4, we write a pair of ordinary differential equations (ODEs) for p and r:

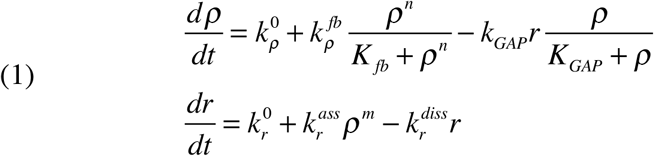

### Estimating model parameter values

We estimated the values for model parameters, we first constructed empirical relationships between intensities of GFP::AHPH and GFP::RGA-3 (herafter RhoA and RGA), and their time derivatives, during pulses of RhoA activity. Intensity data for RhoA and RGA were normalized, aligned and averaged over many individual pulses, then co-aligned using Myosin::RFP as a common reference as described above (Figure 7C). We smoothed these data using a five-frame moving average, and then estimated the time derivatives of signal intensity as the difference in intensities between consecutive frames. We then fit model equations to these data as follows: First, we fixed reference values for basal RhoA activation 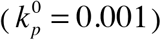 and RGA recruitment (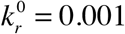), consistent with the slow rates of increase in RhoA and RGA observed before the sharp upswing of each pulse (Figure 7A). To estimate parameters that govern RhoA dynamics, we first considered the rising phase of RhoA activation (colored segments of the curves in Figure S7A), when RGA ~ 0, and used non-linear least squares regression to fit 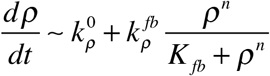 to 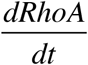 vs *RhoA*. We set n = 1, based on the observed form of dependence of 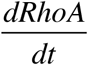 on RhoA (Figure S7D), fixed the value of 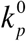, and used model fits to estimate values for 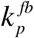 and *K_fb_*. Next, holding values for 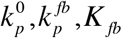 and *n* fixed, we fit the full equation for 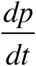 to data for 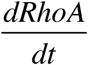 vs. RhoA and RGA for the termination phase (colored segments of the curves in Figure S7B) to estimate values for *k_GAP_* and *K_GAP_*. (Figure 7E). These fits yielded a value for *K_GAP_* that was consistently negative and close to zero, corresponding to a scenario in which RGA-3/4 operates near saturation on active RhoA. Therefore, in subsequent analyses, we set *K_GAP_* to a small constant positive value ( *K_GAP_* = 0.001) and used the model fit to estimate a value for *k_GAP_*. To estimate parameters that govern RhoA dynamics, we fit the full equation for 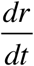 to data for 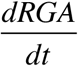 vs RhoA and RGA over the entire pulse.

For each set of parameters determined as above, we set the initial values for r and p to zero, simulating a scenario in which RhoA is minimally active and a small perturbation reduces RGA-3/4 to a minimally observed level. We then solved the equations numerically using Matlab to determine if this initial perturbation would result in either a single pulse of RhoA activity, followed by a return to a stable inactive state (excitability) or a train of pulses (oscillatory dynamics).

## Index of Supplementary Information

### Supplementary Figures

**Figure S1.**Schematic overview of methods for tracking a moving and deforming patch of cortex from single molecule data.

**Figure S2.**Comparison of different methods to quantify local deformation (strain rate) and to align data across multiple pulses.

**Figure S3.**Measurement, alignment and decomposition of fluorescence intensity data during pulsed contractions.

**Figure S4.**Two color analysis of pulsed contractions in P0.

**Figure S5.**Septin localization and contributions to pulsed contractions in one-cell embryos.

### Supplementary Movies

**Movie S1.**Actomyosin pulse dynamics in wild type cells expressing GFP::UTR and NMY-2::RFP.

**Movie S2.**Two cell embryo expressing GFP::ACT-1 at single molecule levels and imaged in near-TIRF mode.

**Movie S3.**Dynamic tracking of a cortical patch from single molecule data.

**Movie S4.**Two cell embryo expressing NMY-2::GFP at single molecule levels and imaged in near-TIRF mode.

**Movie S5.**Pulse dynamics in a two-cell embryo expressing GFP::AHPH and NMY-2::RFP.

**Movie S6.**Pulse dynamics in a zygote P0 expressing GFP::AHPH and NMY-2::mKate and subjected to strong *nmy-2* RNAi.

**Movie S7.**Pulse dynamics in zygotes expressing GFP::AHPH and subjected to *rho-1RNAi* for different periods of time.

**Movie S8.**Dynamics of GFP::RGA-3 and NMY-2::RFP accumulation during pulsing inAB cells.

**Movie S9.**Absence of RhoA pulsing in *rga-3; rga-4* double mutants.

**Movie S10.**Coordinated response of cortical F-actin and RGA-3 to acute treatment with Latrunculin A.

### Author Contributions

Ed Munro conceived the project, and Jon Michaux and Francois Robin shared equally with Ed Munro in its intellectual development. Francois Robin developed and applied the single molecule imaging and particle tracking analysis and performed all the work reported in Figures 2 and 3. Jon Michaux developed the multicolor imaging approach to analysis of pulsed contractions and performed the experiments reported in Figures 4-9. Will McFadden and Ed Munro performed the modeling shown in Figure 10. Jon Michaux, Francois Robin and Ed Munro prepared figures and wrote the paper.

## Acknowledgements

We thank Amy Maddox, Katrina Longhini, Dan Dickinson, Esther Zanin and Karen Oegema for worm strains, Michael Glotzer, Margaret Gardel, Chip Ferguson, and members of the Munro lab for valuable discussions, and Katrina Longini for technical assistance.

This work was supported by NIGMS grants GM09844 (to E.M.M.) and T32GM007197 (to J.B.M.). The content is solely the responsibility of the authors and does not necessarily represent the official views of the National Institutes of Health.

## Abbreviations List

AB: anterior blastomere in the two-cell stage *C. elegans* embryo
AHPH: active RhoA binding domain of Anillin
GAP: GTPase Activating Protein (change on page 5 and in abstract)
NMY-2: non-muscle myosin heavy chain
P0: one-cell *C. elegans* embryo, or zygote
TIRF: total internal reflection (abstract)
UTR: F-actin binding domain of Utrophin

## References

Allard, J., and A. Mogilner. 2013. Traveling waves in actin dynamics and cell motility. Current Opinion in Cell Biology. 25:107–115. doi:10.1016/j.ceb.2012.08.012.

Bement, W.M., M. Leda, A.M. Moe, A.M. Kita, M.E. Larson, A.E. Golding, C. Pfeuti, K.-C. Su, A.L. Miller, A.B. Goryachev, and G. von Dassow. 2015. Activator-inhibitor coupling between Rho signalling and actin assembly makes the cell cortex an excitable medium. Nat. Cell Biol. 17:1471–1483. doi:10.1038/ncb3251.

Blanchard, G.B., S. Murugesu, R.J. Adams, A. Martinez-Arias, and N. Gorfinkiel. 2010. Cytoskeletal dynamics and supracellular organisation of cell shape fluctuations during dorsal closure. Development. 137:2743–2752. doi:10.1242/dev.045872.

Bois, J.S., F. Jülicher, and S.W. Grill. 2011. Pattern formation in active fluids. Phys. Rev. Lett. 106:028103. doi:10.1103/PhysRevLett.106.028103.

Bornens, M., M. Paintrand, and C. Celati. 1989. The cortical microfilament system of lymphoblasts displays a periodic oscillatory activity in the absence of microtubules: implications for cell polarity. The Journal of Cell Biology. 109:1071–1083.

Brenner, S. 1974. The genetics of Caenorhabditis elegans. Genetics. 77:71–94.

Burkel, B.M., G. von Dassow, and W.M. Bement. 2007. Versatile fluorescent probes for actin filaments based on the actin-binding domain of utrophin. Cell Motil. Cytoskeleton. 64:822–832. doi:10.1002/cm.20226.

Carvalho, A., S.K. Olson, E. Gutierrez, K. Zhang, L.B. Noble, E. Zanin, A. Desai, A. Groisman, and K. Oegema. 2011. Acute drug treatment in the early C. elegans embryo. PLoS ONE. 6:e24656. doi:10.1371/journal.pone.0024656.

Chen, Q., S. Nag, and T.D. Pollard. 2012. Formins filter modified actin subunits during processive elongation. J. Struct. Biol. 177:32–39. doi:10.1016/j.jsb.2011.10.005.

Costes, S.V., D. Daelemans, E.H. Cho, Z. Dobbin, G. Pavlakis, and S. Lockett. 2004. Automatic and quantitative measurement of protein-protein colocalization in live cells. Biophys. J. 86:3993–4003. doi:10.1529/biophysj.103.038422.

Crocker, J. 1996. Methods of Digital Video Microscopy for Colloidal Studies. Journal of Colloid and Interface Science. 179:298–310. doi:10.1006/jcis.1996.0217.

David, D.J.V., A. Tishkina, and T.J.C. Harris. 2010. The PAR complex regulates pulsed actomyosin contractions during amnioserosa apical constriction in Drosophila. Development. 137:1645–1655. doi:10.1242/dev.044107.

Dierkes, K., A. Sumi, J. Solon, and G. Salbreux. 2014. Spontaneous oscillations of elastic contractile materials with turnover. Phys. Rev. Lett. 113:148102.

Diogon, M., F. Wissler, S. Quintin, Y. Nagamatsu, S. Sookhareea, F. Landmann, H. Hutter, N. Vitale, and M. Labouesse. 2007. The RhoGAP RGA-2 and LET-502/ROCK achieve a balance of actomyosin-dependent forces in C. elegans epidermis to control morphogenesis. Development. 134:2469–2479. doi:10.1242/dev.005074.

Effler, J.C., Y.-S. Kee, J.M. Berk, M.N. Tran, P.A. Iglesias, and D.N. Robinson. 2006. Mitosis-specific mechanosensing and contractile-protein redistribution control cell shape. Current Biology. 16:1962–1967. doi:10.1016/j.cub.2006.08.027.

Fernandez-Gonzalez, R., S. de M. Simões, J.-C. Röper, S. Eaton, and J.A. Zallen. 2009. Myosin II dynamics are regulated by tension in intercalating cells. Dev. Cell. 17:736–743. doi:10.1016/j.devcel.2009.09.003.

Goldbeter, A. 1996. Biochemical Oscillations and Cellular Rhythms: The Molecular Bases of Periodic and Chaotic Behavior. Cambridge Univ. Press, Cambridge, UK.

Gorfinkiel, N. 2016. From actomyosin oscillations to tissue-level deformations. Dev Dyn. 245:268–275. doi:10.1002/dvdy.24363.

Graessl, M., J. Koch, A. Calderon, D. Kamps, S. Banerjee, T. Mazel, N. Schulze, J.K. Jungkurth, R. Patwardhan, D. Solouk, N. Hampe, B. Hoffmann, L. Dehmelt, and P. Nalbant. 2017. An excitable Rho GTPase signaling network generates dynamic subcellular contraction patterns. The Journal of Cell Biology. 216:4271–4285. doi:10.1083/jcb.201706052.

Hamill, D.R., A.F. Severson, J.C. Carter, and B. Bowerman. 2002. Centrosome maturation and mitotic spindle assembly in C. elegans require SPD-5, a protein with multiple coiled-coil domains. Dev. Cell. 3:673–684.

Hayakawa, K., H. Tatsumi, and M. Sokabe. 2011. Actin filaments function as a tension sensor by tension-dependent binding of cofilin to the filament. The Journal of Cell Biology. 195:721–727. doi:10.1038/31735.

He, L., X. Wang, H.L. Tang, and D.J. Montell. 2010. Tissue elongation requires oscillating contractions of a basal actomyosin network. Nat. Cell Biol. 12:1133–1142. doi:10.1038/ncb2124.

Hickson, G.R.X., and P.H. O’Farrell. 2008. Rho-dependent control of anillin behavior during cytokinesis. The Journal of Cell Biology. 180:285–294. doi:10.1083/jcb.200709005.

Izhikevich, E.M. 2007. Dynamical systems in neuroscience : the geometry of excitability and bursting. MIT Press, Cambridge, Mass., London.

Jaffe, A.B., and A. Hall. 2005. Rho GTPases: biochemistry and biology. Annu Rev Cell Dev Biol. 21:247–269. doi:10.1146/annurev.cellbio.21.020604.150721.

Kamath, R.S., A.G. Fraser, Y. Dong, G. Poulin, R. Durbin, M. Gotta, A. Kanapin, N. Le Bot, S. Moreno, M. Sohrmann, D.P. Welchman, P. Zipperlen, and J. Ahringer. 2003. Systematic functional analysis of the Caenorhabditis elegans genome using RNAi. Nature. 421:231–237. doi:10.1038/nature01278.

Kapustina, M., G.E. Weinreb, N. Costigliola, Z. Rajfur, K. Jacobson, and T.C. Elston. 2008. Mechanical and biochemical modeling of cortical oscillations in spreading cells. Biophys. J. 94:4605–4620. doi:10.1529/biophysj.107.121335.

Kapustina, M., T.C. Elston, and K. Jacobson. 2013. Compression and dilation of the membrane-cortex layer generates rapid changes in cell shape. The Journal of Cell Biology. 200:95–108. doi:10.1083/jcb.201204157.

Kasza, K.E., and J.A. Zallen. 2011. Dynamics and regulation of contractile actin-myosin networks in morphogenesis. Current Opinion in Cell Biology. 23:30–38. doi:10.1016/j.ceb.2010.10.014.

Kim, H.Y., and L.A. Davidson. 2011. Punctuated actin contractions during convergent extension and their permissive regulation by the non-canonical Wnt-signaling pathway. J Cell Sci. 124:635–646. doi:10.1242/jcs.067579.

Koride, S., L. He, L.-P. Xiong, G. Lan, D.J. Montell, and S.X. Sun. 2014. Mechanochemical regulation of oscillatory follicle cell dynamics in the developing Drosophila egg chamber. Mol Biol Cell. 25:3709–3716. doi:10.1091/mbc.E14-04-0875.

Kumar, K.V., J.S. Bois, F. Jülicher, and S.W. Grill. 2014. Pulsatory Patterns in Active Fluids. Phys. Rev. Lett. 112:208101. doi:10.1103/PhysRevLett.112.208101.

La Cruz De, E.M., and M.L. Gardel. 2015. Actinn, mechanics and fragmentation. Journal of Biological Chemistry. jbc. R115.636472. doi:10.1074/jbc.R115.636472.

Landau, L.D., and E.M. Lifshitz. 1987. Fluid Mechanics, Second Edition: nn Volume 6 (Course of Theoretical Physics). Butterworth-Heinemann.

Levayer, R., and T. Lecuit. 2012. Biomechanical regulation of contractility: spatial control and dynamics. Trends Cell Biol. 22:61–81. doi:10.1016/j.tcb.2011.10.001.

Levayer, R., and T. Lecuit. 2013. Oscillation and polarity of E-cadherin asymmetries control actomyosin flow patterns during morphogenesis. Dev. Cell. 26:162–175. doi:10.1016/j.devcel.2013.06.020.

Loria, A., K.M. Longhini, and M. Glotzer. 2012. The RhoGAP domain of CYK-4 has an essential role in RhoA activation. Curr Biol. 22:213–219. doi:10.1016/j.cub.2011.12.019.

Luo, C.H., and Y. Rudy. 1991. A model of the ventricular cardiac action potential. Depolarization, repolarization, and their interaction. Circulation Research. 68:1501–1526.

Luo, T., K. Mohan, V. Srivastava, Y. Ren, P.A. Iglesias, and D.N. Robinson. 2012. Understanding the cooperative interaction between myosin II and actin cross-linkers mediated by actin filaments during mechanosensation. Biophys. J. 102:238–247. doi:10.1016/j.bpj.2011.12.020.

Machado, P.F., G.B. Blanchard, J. Duque, and N. Gorfinkiel. 2014. Cytoskeletal turnover and Myosin contractility drive cell autonomous oscillations in a model of Drosophila Dorsal Closure. Eur. Phys. J. Spec. Top. 223:1391–1402. doi:10.1140/epjst/e2014-02197-7.

Maddox, A.S., B. Habermann, A. Desai, and K. Oegema. 2005. Distinct roles for two C. elegans anillins in the gonad and early embryo. Development. 132:2837–2848. doi:10.1242/dev.01828.

Maddox, A.S., L. Lewellyn, A. Desai, and K. Oegema. 2007. Anillin and the septins promote asymmetric ingression of the cytokinetic furrow. Dev. Cell. 12:827–835. doi:10.1016/j.devcel.2007.02.018.

Martin, A.C., M. Kaschube, and E.F. Wieschaus. 2009. Pulsed contractions of an actin-myosin network drive apical constriction. Nature. 457:495–499. doi:10.1038/nature07522.

Mason, F.M., S. Xie, C.G. Vasquez, M. Tworoger, and A.C. Martin. 2016. RhoA GTPase inhibition organizes contraction during epithelial morphogenesis. The Journal of Cell Biology. 214:603–617. doi:10.1083/jcb.201603077.

Motegi, F., and A. Sugimoto. 2006. Sequential functioning of the ECT-2 RhoGEF, RHO-1 and CDC-42 establishes cell polarity in Caenorhabditis elegans embryos. Nat. Cell Biol. 8:978–985. doi:10.1038/ncb1459.

Munjal, A., J.-M. Philippe, E. Munro, and T. Lecuit. 2015. A self-organized biomechanical network drives shape changes during tissue morphogenesis. Nature. 524:351–355. doi:10.1038/nature14603.

Munro, E., J. Nance, and J.R. Priess. 2004. Cortical flows powered by asymmetrical contraction transport PAR proteins to establish and maintain anterior-posterior polarity in the early C. elegans embryo. Dev. Cell. 7:413–424. doi:10.1016/j.devcel.2004.08.001.

Naganathan, S.R., S. Fürthauer, M. Nishikawa, F. Jülicher, and S.W. Grill. 2014. Active torque generation by the actomyosin cell cortex drives left-right symmetry breaking. Elife. 3:e04165. doi:10.7554/eLife.04165.

Nishikawa, M., S.R. Naganathan, F. Jülicher, and S.W. Grill. 2017. Controlling contractile instabilities in the actomyosin cortex. Elife. 6:058101. doi:10.7554/eLife.19595.

Olson, S.K., G. Greenan, A. Desai, T. Müller-Reichert, and K. Oegema. 2012. Hierarchical assembly of the eggshell and permeability barrier in C. elegans. The Journal of Cell Biology. 198:731–748. doi:10.1083/jcb.201206008.

Paluch, E., M. Piel, J. Prost, M. Bornens, and C. Sykes. 2005. Cortical actomyosin breakage triggers shape oscillations in cells and cell fragments. Biophys. J. 89:724–733. doi:10.1529/biophysj.105.060590.

Pelletier, V., N. Gal, P. Fournier, and M.L. Kilfoil. 2009. Microrheology of Microtubule Solutions and Actin-Microtubule Composite Networks. Phys. Rev. Lett. 102:188303. doi:10.1103/PhysRevLett.102.188303.

Piekny, A.J., and A.S. Maddox. 2010. The myriad roles of Anillin during cytokinesis. Semin. Cell Dev. Biol. 21:881–891. doi:10.1016/j.semcdb.2010.08.002.

Piekny, A.J., and M. Glotzer. 2008. Anillin is a scaffold protein that links RhoA, actin, and myosin during cytokinesis. Current Biology. 18:30–36. doi:10.1016/j.cub.2007.11.068.

Piekny, A.J., and P.E. Mains. 2002. Rho-binding kinase (LET-502) and myosin phosphatase (MEL-11) regulate cytokinesis in the early Caenorhabditis elegans embryo. J Cell Sci. 115:2271–2282.

Piekny, A.J., J.-L.F. Johnson, G.D. Cham, and P.E. Mains. 2003. The Caenorhabditis elegans nonmuscle myosin genes nmy-1 and nmy-2 function as redundant components of the let-502/Rho-binding kinase and mel-11/myosin phosphatase pathway during embryonic morphogenesis. Development. 130:5695–5704. doi:10.1242/dev.00807.

Pohl, C., M. Tiongson, J.L. Moore, A. Santella, and Z. Bao. 2012. Actomyosin-based self-organization of cell internalization during C. elegans gastrulation. BMC Biol. 10:94. doi:10.1186/1741-7007-10-94.

Rankin, K.E., and L. Wordeman. 2010. Long astral microtubules uncouple mitotic spindles from the cytokinetic furrow. The Journal of Cell Biology. 190:35–43. doi:10.1083/jcb.201004017.

Rauzi, M., P.-F. Lenne, and T. Lecuit. 2010. Planar polarized actomyosin contractile flows control epithelial junction remodelling. Nature. 468:1110–1114. doi:10.1038/nature09566.

Razzell, W., W. Wood, and P. Martin. 2014. Recapitulation of morphogenetic cell shape changes enables wound re-epithelialisation. Development. 141:1814–1820. doi:10.1242/dev.107045.

Ren, Y., J.C. Effler, M. Norstrom, T. Luo, R.A. Firtel, P.A. Iglesias, R.S. Rock, and D.N. Robinson. 2009. Mechanosensing through cooperative interactions between myosin II and the actin crosslinker cortexillin I. Curr Biol. 19:1421–1428. doi:10.1016/j.cub.2009.07.018.

Robin, F.B., W.M. McFadden, B. Yao, and E.M. Munro. 2014. Single-molecule analysis of cell surface dynamics in Caenorhabditis elegans embryos. Nat Meth. 11:677–682. doi:10.1038/nmeth.2928.

Salbreux, G., J.F. Joanny, J. Prost, and P. Pullarkat. 2007. Shape oscillations of non-adhering fibroblast cells. Phys Biol. 4:268–284. doi:10.1088/1478-3975/4/4/004.

Schiffhauer, E.S., T. Luo, K. Mohan, V. Srivastava, X. Qian, E.R. Griffis, P.A. Iglesias, and D.N. Robinson. 2016. Mechanoaccumulative Elements of the Mammalian Actin Cytoskeleton. Curr Biol. 26:1473–1479. doi:10.1016/j.cub.2016.04.007.

Schmutz, C., J. Stevens, and A. Spang. 2007. Functions of the novel RhoGAP proteins RGA-3 and RGA-4 in the germ line and in the early embryo of C. elegans. 134:3495–3505. doi:10.1242/dev.000802.

Schonegg, S., A.T. Constantinescu, C. Hoege, and A.A. Hyman. 2007. The Rho GTPase-activating proteins RGA-3 and RGA-4 are required to set the initial size of PAR domains in Caenorhabditis elegans one-cell embryos. 104:14976–14981. doi:10.1073/pnas.0706941104.

Schonegg, S., and A.A. Hyman. 2006. CDC-42 and RHO-1 coordinate acto-myosin contractility and PAR protein localization during polarity establishment in C. elegans embryos. 133:3507–3516. doi:10.1242/dev.02527.

Sedzinski, J., M. Biro, A. Oswald, J.-Y. Tinevez, G. Salbreux, and E. Paluch. 2011. Polar actomyosin contractility destabilizes the position of the cytokinetic furrow. Nature. 476:462–466. doi:10.1038/nature10286.

Segal, D., A. Zaritsky, E.D. Schejter, and B.-Z. Shilo. 2018. Feedback inhibition of actin on Rho mediates content release from large secretory vesicles. The Journal of Cell Biology. 217:1815–1826. doi:10.1083/jcb.201711006.

Severson, A.F., D.L. Baillie, and B. Bowerman. 2002. A Formin Homology protein and a profilin are required for cytokinesis and Arp2/3-independent assembly of cortical microfilaments in C. elegans. Current Biology. 12:2066–2075.

Solon, J., A. Kaya-Copur, J. Colombelli, and D. Brunner. 2009. Pulsed forces timed by a ratchet-like mechanism drive directed tissue movement during dorsal closure. Cell. 137:1331–1342. doi:10.1016/j.cell.2009.03.050.

Strogatz, S. 1994. Nonlinear dynamics and chaos with applications to physics, biology, chemistry, and engineering. amazon.ca.

Timmons, L., D.L. Court, and A. Fire. 2001. Ingestion of bacterially expressed dsRNAs can produce specific and potent genetic interference in Caenorhabditis elegans. Gene. 263:103–112.

Tokunaga, M., N. Imamoto, and K. Sakata-Sogawa. 2008. Highly inclined thin illumination enables clear single-molecule imaging in cells. Nat Meth. 5:159–161. doi:10.1038/nmeth1171.

Tse, Y.C., A. Piekny, and M. Glotzer. 2011. Anillin promotes astral microtubule-directed cortical myosin polarization. Mol Biol Cell. 22:3165–3175. doi:10.1091/mbc.E11-05-0399.

Tse, Y.C., M. Werner, K.M. Longhini, J.-C. Labbé, B. Goldstein, and M. Glotzer. 2012. RhoA activation during polarization and cytokinesis of the early Caenorhabditis elegans embryo is differentially dependent on NOP-1 and CYK-4. Mol Biol Cell. 23:4020–4031. doi:10.1091/mbc.E12-04-0268.

Vallotton, P., S.L. Gupton, C.M. Waterman-Storer, and G. Danuser. 2004. Simultaneous mapping of filamentous actin flow and turnover in migrating cells by quantitative fluorescent speckle microscopy. PNAS. 101:9660–9665. doi:10.1073/pnas.0300552101.

Vasquez, C.G., M. Tworoger, and A.C. Martin. 2014. Dynamic myosin phosphorylation regulates contractile pulses and tissue integrity during epithelial morphogenesis. The Journal of Cell Biology. 206:435–450. doi:10.1083/jcb.201402004.

Watanabe, N., and T.J. Mitchison. 2002. Single-molecule speckle analysis of actin filament turnover in lamellipodia. Science. 295:1083–1086. doi:10.1126/science.1067470.

Weiner, O.D., W.A. Marganski, L.F. Wu, S.J. Altschuler, and M.W. Kirschner. 2007. An actin-based wave generator organizes cell motility. PLoS Biol. 5:e221. doi:10.1371/journal.pbio.0050221.

Werner, M., E. Munro, and M. Glotzer. 2007. Astral signals spatially bias cortical myosin recruitment to break symmetry and promote cytokinesis. Current Biology. 17:1286–1297. doi:10.1016/j.cub.2007.06.070.

Zanin, E., A. Desai, I. Poser, Y. Toyoda, C. Andree, C. Moebius, M. Bickle, B. Conradt, A. Piekny, and K. Oegema. 2013. A conserved RhoGAP limits M phase contractility and coordinates with microtubule asters to confine RhoA during cytokinesis. Dev. Cell. 26:496–510. doi:10.1016/j.devcel.2013.08.005.

Zhang, D., and M. Glotzer. 2015. The RhoGAP activity of CYK-4/MgcRacGAP functions non-canonically by promoting RhoA activation during cytokinesis. Elife. 4:204. doi:10.7554/eLife.08898.

